# AbDesign: Database of point mutants of antibodies with associated structures reveals poor generalization of binding predictions from machine learning models

**DOI:** 10.1101/2025.06.09.658639

**Authors:** Bartosz Janusz, Dawid Chomicz, Samuel Demharter, Marloes Arts, Jurrian de Kanter, Yano Wilke, Helena Britze, Sonia Wrobel, Tomasz Gawłowski, Pawel Dudzic, Kärt Ukkivi, Lauri Peil, Roberto Spreafico, Konrad Krawczyk

## Abstract

Antibodies are naturally evolved molecular recognition scaffolds that can bind a variety of surfaces. Their designability is crucial to the development of biologics, with computational methods holding promise in accelerating the delivery of medicines to the clinic. Modeling antibody-antigen recognition is prohibitively difficult, with data paucity being one of the biggest hurdles. Current affinity datasets comprise a small number of experimental measurements, which are often not standardized between molecules. Here, we address these issues by creating a dataset of seven antigens with two antibodies each, for which we introduce a heterogeneous set of mutations to the CDR-H3 measured by ELISA. Each of the parental complexes has a known crystal structure. We perform benchmarking of state-of-the-art affinity prediction algorithms to gauge their effectiveness. Current computational methods exhibit significant limitations in accurately predicting the effects of single-point mutations. In contrast, the older empirical, physics-based method FoldX, performs well in identifying mutants that retain binding. These findings highlight the need for more resources like the one presented here — large, molecularly diverse, and experimentally consistent datasets.

## Introduction

Antibodies are among the most critical classes of biologics, serving as pivotal tools in diagnostics, therapeutics, and research ^1^. Their ability to specifically bind to a vast array of antigens makes them invaluable in treating diseases such as cancer, autoimmune disorders, and infectious diseases. Given the significance of biologics, the development of computational methods to predict and enhance antibody affinity is of paramount importance^2,3^. It remains an ambition of the biologics drug discovery community for such methods to accelerate the design and optimization of antibodies, reducing the time and cost associated with experimental approaches. In practice, such efforts are frustrated by the paucity of available data.

Structural information plays a crucial role in the design and development of novel antibody binders ^4^. The availability of high-resolution structures of antibodies bound to their antigens provides detailed insights into the molecular interactions that govern binding affinity and specificity. Resources such as the Protein Data Bank (PDB^5^) serve as a primary repository for these structures, offering a wealth of data for computational analyses. Databases such as the Structural Antibody Database (SAbDab^6^) and the Antibody Structure Database (AbDb^7^) curate and organize this structural information, enabling researchers to efficiently retrieve and analyze antibody-antigen complexes.

These databases are instrumental in informing the design of new binders by providing templates for modeling, docking simulations, and structure-based predictions of antibody affinity.

While structural information is indispensable, it does not provide any information about the strength of the interaction. The creation of computational methods for developing antibodies is heavily reliant on the availability of robust affinity databases. Databases such as SKEMPI 2.0^8^ and AB-Bind^9^ are central to this effort, as they provide curated collections of experimental data on the affinities of antibody-antigen complexes. Though SKEMPI focuses more broadly on the effects of mutations on protein-protein interaction affinities, antibodies form a subset of the data. Similarly, the AB-Bind database provides detailed information on the binding affinities of antibodies, allowing for the development of predictive models that can guide the design of high-affinity binders. These databases serve as critical resources for the development of machine learning algorithms and other computational tools aimed at predicting and optimizing antibody interactions.

Most recently, an affinity dataset accompanying IgDesign has been released, comprising ca. 1,000 variants of antibody therapeutics for seven targets^10^. While this is a very useful dataset, many of the mutants show a large edit distance from the parental antibody for which a structure is available. Since antibody-antigen modeling is still not a solved problem^11^, it is unfeasible to obtain reliable 3D conformations of variants to model affinity measurements in their structural context.

Despite the progress in developing computational methods for antibody affinity prediction, several challenges persist due to limitations in the current experimental datasets. One significant issue is the relatively small size of these resources, which constrains the statistical power of computational models. There are approximately 1,000 antibody-antigen affinity data points, including single as well as higher-order mutations, between SKEMPI and AB-Bind. Additionally, the experimental methods used to generate affinity data are not uniform, leading to inconsistencies and difficulties in comparing results across different studies. There is also a notable bias towards mutations of a particular kind, such as alanine-scanning, which limits the ability of models to generalize to more complex modification scenarios. More plentiful binder datasets lack corresponding structural data^12–14^, hindering the ability to incorporate conformational insights into predictive models. These challenges underscore the need for more comprehensive and standardized experimental datasets to support the continued advancement of computational methods in antibody development.

Here we present the AbDesign database, a resource of 14 complexes (7 antigens with two antibodies each, with publicly available structures), to which heterogeneous point mutations were introduced to the binding residues of the CDR-H3. Since there are ca. 650 antibody point mutants between SKEMPI and AB-Bind, our contribution of 672 points increases the amount of available single-mutant data twofold and avoids common biases, such as predominant alanine-scanning.

## Materials and Methods

### Selection of Antigens/Antibodies and Mutation Strategy

We aimed to select antibody mutants covering seven plates with 96 variants each. Each plate contained antibody variants binding to a single target antigen. To introduce diversity, for each of the seven target antigens we selected two distinct parental antibodies. According to these criteria, we identified seven target antigens from the PDB, each with two structurally distinct antibodies available (Table 1). While targets were chosen for their presence in therapeutic or frequently studied contexts (e.g., PD-1, CTLA-4, VEGF, etc.), our primary requirement was to ensure at least one crystal structure per antibody-antigen pair. The structures were selected on the basis of the quality of the model and greater resolutions. This step was carried out manually since it involved a selection of just 14 structures.

**Table 1.**
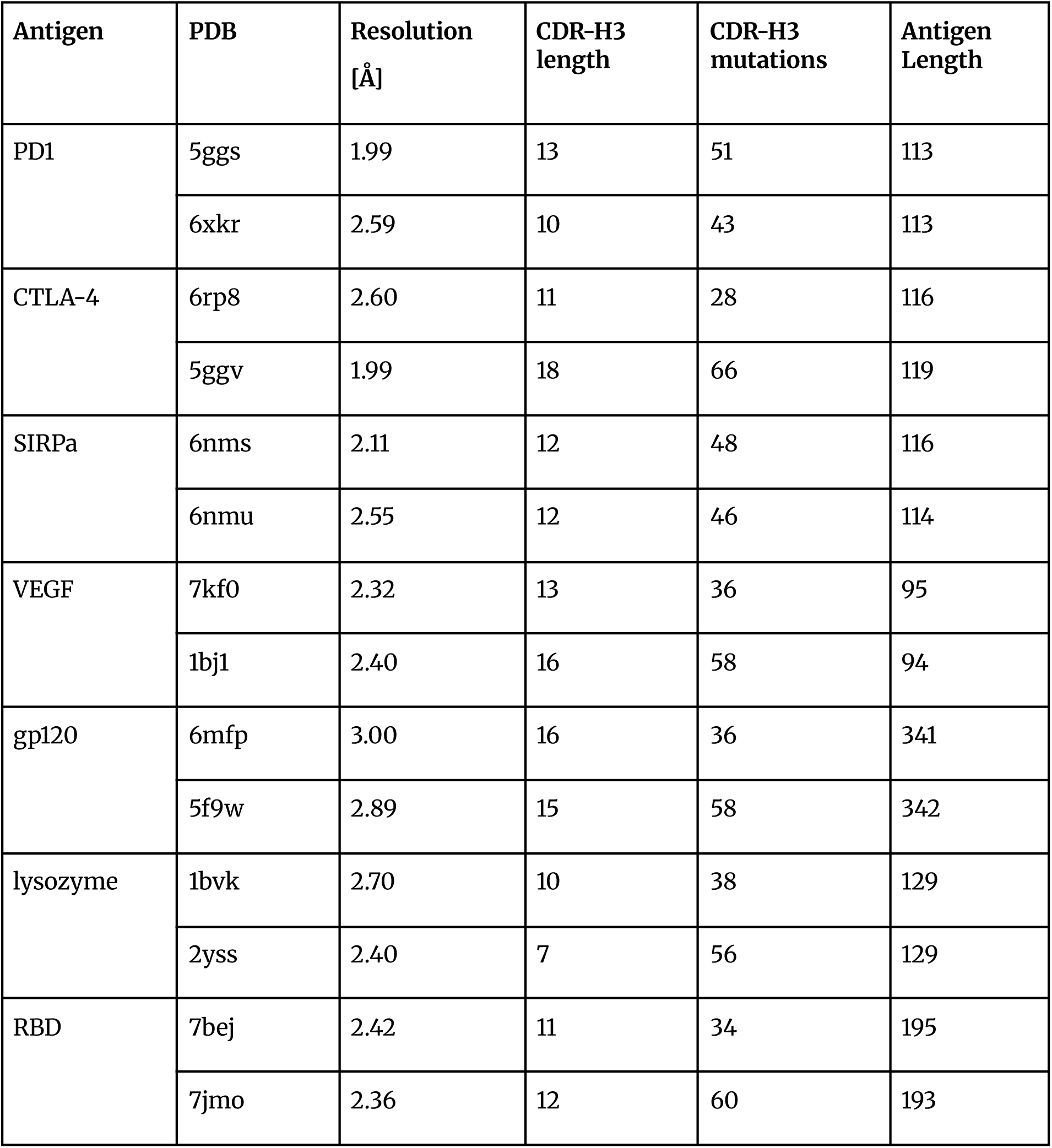
Contents of the AbDesign Database. We selected fourteen PDB structures of antibodies in complexes with seven antigens, i.e., two antibodies per antigen, to introduce mutations. We only modified the positions of CDR-H3 less than 4.5Å away from the antigen, based on heavy atom distance. We aimed for a uniform number of mutations spread across positions within each antibody.

| Antigen | PDB | Resolution [Å] | CDR-H3 length | CDR-H3 mutations | Antigen Length |
| --- | --- | --- | --- | --- | --- |
| PD1 | 5ggs | 1.99 | 13 | 51 | 113 |
|  | 6xkr | 2.59 | 10 | 43 | 113 |
| CTLA-4 | 6rp8 | 2.60 | 11 | 28 | 116 |
|  | 5ggv | 1.99 | 18 | 66 | 119 |
| SIRPa | 6nms | 2.11 | 12 | 48 | 116 |
|  | 6nmu | 2.55 | 12 | 46 | 114 |
| VEGF | 7kf0 | 2.32 | 13 | 36 | 95 |
|  | 1bj1 | 2.40 | 16 | 58 | 94 |
| gp120 | 6mfp | 3.00 | 16 | 36 | 341 |
|  | 5f9w | 2.89 | 15 | 58 | 342 |
| lysozyme | 1bvk | 2.70 | 10 | 38 | 129 |
|  | 2yss | 2.40 | 7 | 56 | 129 |
| RBD | 7bej | 2.42 | 11 | 34 | 195 |
|  | 7jmo | 2.36 | 12 | 60 | 193 |

We focused specifically on CDR-H3 residues that were within 4.5 Å of the antigen’s heavy atoms, as identified by computational contact mapping. Each residue in that region was systematically mutated to create a comprehensive panel of 658 single-point mutants across the 14 complexes, plus 14 wild-type (parental) measurements (one per complex, as controls), totalling 672 variants. As the number of contacting residues was not uniform across antibodies, and each plate could only house 96 variants, a sampling scheme was introduced to choose mutations uniformly (excluding cysteine). We aimed to select as many of the 18 mutant amino acids for each position as possible. In cases where this was impossible, the selection of variants was adjusted manually, so as to maintain the heterogeneity of the dataset. However, due to the available plate budget, it was unfeasible to systematically cover all possible mutations at each position, leading to unequal sampling of different amino acids. Still, sampling bias was lower than that found in any other antibody-antigen dataset, as discussed in the Results section.

### Antibody Production and Purification

Antibodies were produced by Icosagen using its QMCF Technology platform. For all antibodies, VH and VL inserts were cloned into pQMCF bi-cistronic expression vectors, containing human IgG1 constant region and human kappa light chain constant region. Sequence-verified expression vectors were then chemically transfected with reagent R007 into CHOEBNALT-1E9 cells. Cells were cultivated in CHO TF Medium (Sartorius, Cat# 886-0001) on 96-deepwell plates. At the end of the production, cells were removed by centrifugation, and antibodies were purified from the cell culture supernatant with Mag Sepharose™ PrismA magnetic beads (Cytiva, Cat #17550001), using the KingFisher Flex purification system (Thermo Scientific, Cat# 711-8G0777). Purified antibodies were formulated into 1x PBS pH 7.4 buffer using Zeba™ Spin Desalting Plates (Thermo Scientific, Cat #87775).

### ELISA Measurements

All measurements were performed using identical ELISA protocols across the entire dataset to ensure direct comparability.

Antigens (Table 2) were diluted in PBS pH 7.4 and added to Clear Flat-Bottom MaxiSorp 96-well plates (Thermo Scientific) at concentrations of 1 µg/mL. After incubation at 4°C for 16-20h and 4x washing with PBS, 0.05 % Tween 20, 0.1% Proclin 300 buffer, wells were blocked for 1 h at room temperature with 1x PBS, 2% BSA, 0.05% Tween20, 0.2% Proclin 300 buffer. After 4x washing, each antibody variant was added to the plate in 4-fold serial dilutions starting from 1 µg/mL in 1x PBS, 1% BSA, 0.05% Tween20, 0.2% Proclin 300 buffer, and incubated at room temperature for 1 h. After 4x washing, Goat anti-Human IgG Fc Cross-Adsorbed Secondary Antibody (Invitrogen, Cat# A18823) was added to the wells at a 1:10 000 dilution and incubated at room temperature for 1 h. After 4x washing, TMB VII substrate was added to the wells, and the reaction was stopped by adding 0.5 M H₂SO₄. Absorbance was measured at 450 nm using Thermo Scientific Multiskan Fc.

**Table 2.** Antigens used in the study.

| Antigen | Supplier |
| --- | --- |
| SARS-CoV-2 RBD2 | Icosagen, Cat# P-307-100 |
| Lysozyme antigen | Creative Diagnostics, Cat# DAG-T1235 |
| HIV-1 GP120 protein, His tag | AcroBiosystems, Cat# GP4-V15223 |
| Human VEGF-165 Recombinant Protein | Gibco, Cat# PHC9391 |
| SIRP alpha Protein, Human, Recombinant (ECD, SIRP alphaV2, His Tag) | SinoBiologicals, Cat# 30014-H08H |
| Human CTLA-4 His-tag Recombinant Protein | Invitrogen, Cat# A42594 |
| Human PD-1 His-tag Recombinant Protein | Invitrogen; Cat# A42533 |

Each measurement was normalized to the absorbance of the wild-type antibody measured on the same plate, yielding a ratio >1 or <1, which captures whether the mutant showed stronger or weaker binding than parental.

### Structural Modeling

Crystal structures for every wild-type complex specified in the three datasets (SKEMPI, AB-Bind, AbDesign) were downloaded from the RCSB Protein Data Bank. All PDB models were cleaned (removing hydrogens, alternative location atoms, and renumbering from 1 to n), antibody chains trimmed to variable regions using RIOT^15^ and correct triplets of heavy-light-antigen chains were manually selected. We treated the resulting cleaned PDB files as a starting point for two structural modeling pipelines:

ABodyBuilder2-based^16^ and conformation-based. ABodyBuilder2 is a neural network that predicts the coordinates of the antibody sequence and we merged the resulting mutated model with the clean structure. In case of conformational sampling, the most common side-chain was replaced in the cleaned structure.

### ABodyBuilder2 Structure Prediction

We used ABodyBuilder2 (ABB2) to model structures of WT antibodies as well as every single position mutant. ABB2 uses a pair of heavy and light chain sequences as input. They are fed into four distinct deep-learning models to generate a diverse set of structural predictions. The structure closest to the ensemble’s average is then chosen. Figure 1 shows an example of ABB2 prediction results.

**Figure 1.**
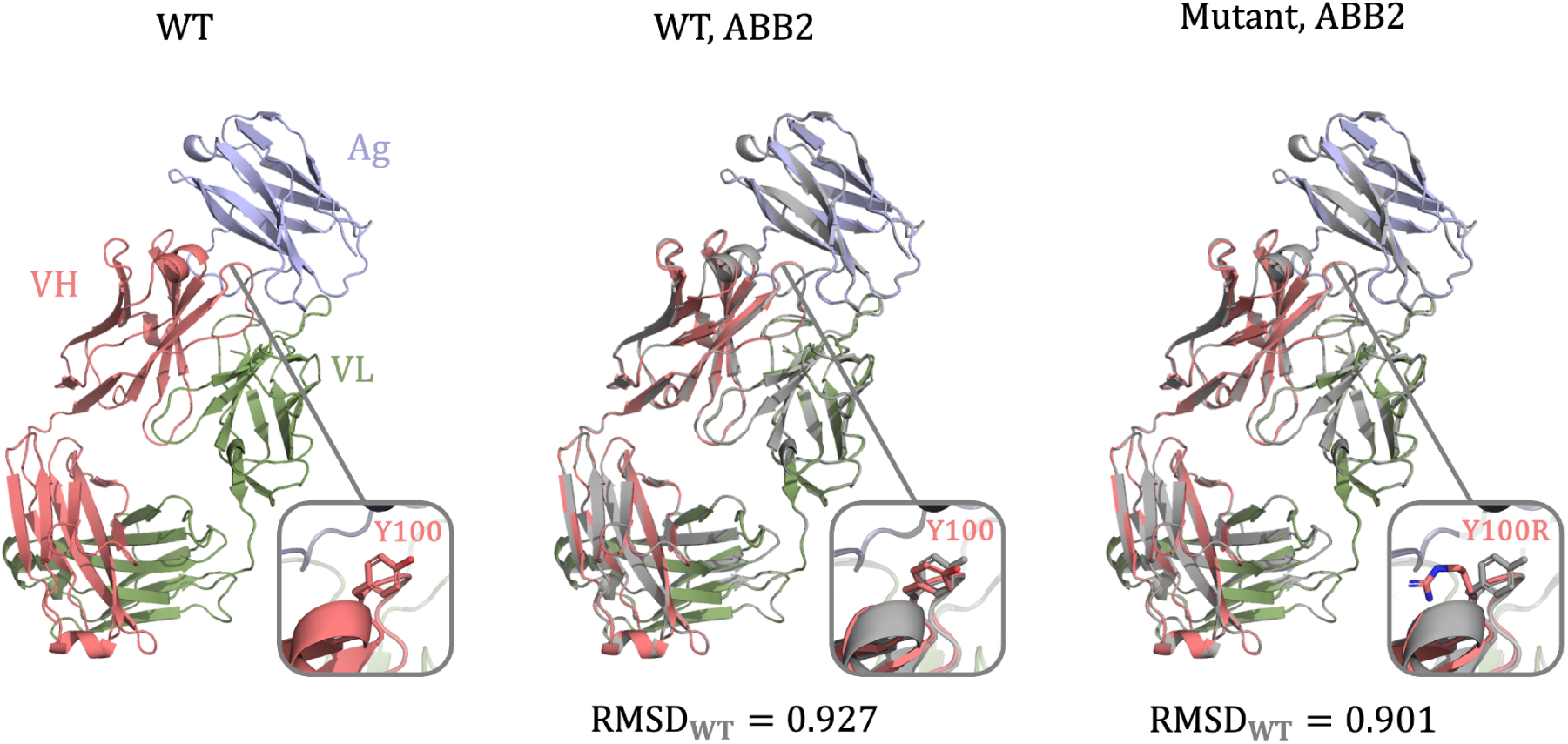
ABodyBuilder2 structure prediction. **A.** Wild-type experimental structure of 5GGS, with tyrosine at position 100. **B.** Wild-type structure predicted by ABB2 with RMSD calculated against experimental wild-type structure. **C.** Mutant structure predicted by ABB2, where the tyrosine at position 100 was mutated to an arginine.

#### Structure Refinement

The neural network within ABodyBuilder2 can sometimes produce physically implausible artifacts, such as steric clashes, incorrect bond lengths, or unwanted cis-peptide bonds, which prompted the authors to develop post-processing steps to address these. To correct these issues, OpenMM with the AMBER14 force field was used for energy minimization, followed by pdbfixer to resolve severe steric clashes by remodeling side chains ^17^. Additional constraints were used to ensure peptide bonds adopt the trans configuration. To prevent L-amino acids from flipping into their D-forms, hydrogen positions at the alpha carbon were adjusted before final relaxation.

To carry out the steps just described, we used the *refine* function from ABB2 with all default parameters, except for *check_for_strained_bonds*, which was set to *False.* This was motivated by the fact that the default *True* setting would lead to trimming of some residues from chain endings‒an undesirable outcome given that all ΔΔG prediction models require WT and mutant structures to have the same length‒thereby causing errors during prediction.

#### Docking predicted antibody structure with antigen

ABB2 models only the antibody part of the complex. To maintain the proper structure of the entire complex, we align the framework residues of the ABB2-generated models with the original crystal structure of the WT. Next, we remove the crystal structure, leaving predicted antibody and crystal antigen structures docked.

### Conformation-based Structure Prediction

The second mutation structure prediction pipeline does not involve any deep learning models, but rather leverages sampling from a rotamer library. We take WT crystal structures and, for the specified single-point mutation, we replace the original residue side-chain with the mutant one. We choose the side-chain atom coordinates from the set of possible conformations, selecting the cluster representative with the highest cardinality and no detected clashes with the crystal structure. After the substitution, both WT and mutated structure are processed using ABB2’s *refine* function, described in the previous paragraph.

### Clustering Procedure and Representatives Selection

We aimed to build a comprehensive database of observed side-chain conformations at each residue position. First, all antibody structures from the RCSB database are collected, renumbered according to IMGT numbering, and deduplicated using sequence identity-based clustering. This step ensures that redundant structures do not bias subsequent analysis. For each IMGT position across all structures, and for each of the 20 standard amino acid types, side-chain atomic coordinates are extracted and aligned based on the backbone atoms. To quantify the structural diversity of side-chain conformations, pairwise root-mean-square deviation (RMSD) values are calculated between all side-chain atom configurations for a given (residue type, position) pair. These RMSD distances serve as input for agglomerative clustering, allowing representative conformations to be selected for each group. Finally, representative side-chain conformations are filtered to ensure that they do not introduce steric clashes with the corresponding backbone context. The resulting database of residue-position-dependent conformations is later used in the conformation modeling pipeline to propose realistic side-chain placements for mutated positions during antibody design.

### Final Structural Dataset

The resulting set of 3D structures includes multiple variants for structural and computational analysis. This includes wild-type crystal structures, both in their original form and refined using ABB2’s *refine* procedure. Additionally, the dataset contains ABB2-generated wild-type structures that have been refined, as well as single-point mutated variants produced through side-chain replacement and subsequent refinement. These models are optimized for direct application in structure-based energy calculations and machine learning workflows.

### Benchmarking Affinity Models

Three modern machine learning models were selected for benchmarking, based on their reported performance: DSMBind, Binding-DDG-predictor, and RDE-PPI (Table 3). To compare with a well-established empirical method, we sourced pre-calculated FoldX datapoints published by Hummer et al.^18^.

**Table 3.** Machine learning models employed for benchmarking as part of this study.

| Model | Inputs | Outputs | Dataset & training | Reference |
| --- | --- | --- | --- | --- |
| **DSMBind** | WT PDB, mutated PDB | $\Delta G$ separately for WT and mutated | SKEMPI 2.0, training on all PPI data and fine-tuning on antibody-antigen complexes. | <sup>19</sup> |
| **Binding-DDG-predictor** | WT PDB, mutated PDB | $\Delta\Delta G$ (mutant - WT) | SKEMPI 2.0, split by complex, five-fold cross validation. | <sup>20</sup> |
| **RDE-PPI** | WT PDB, mutation instructions (yaml) | $\Delta\Delta G$ (mutant - WT) | SKEMPI 2.0, three-fold cross-validation. | <sup>21</sup> |

DSMBind is an unsupervised energy-based model that predicts binding affinities via SE(3) denoising score matching, leveraging unlabeled crystal structures to infer binding energy. While its core framework is unsupervised, supervised variants have been developed for specific tasks, including a version fine-tuned on antibody-antigen interactions. This variant combines DSMBind’s SE(3)-equivariant FANN encoder with ESM-2 sequence embeddings, pre-trained on SKEMPI’s protein-protein binding data and refined on its antibody-antigen subset. It achieves competitive performance (Spearman ⍴ = 0.35) in antibody binding prediction, surpassing methods like AlphaFold2 and protein language models. We used this supervised variant in our benchmarks.

The Binding-DDG-predictor employs an attention-based geometric neural network to predict changes in binding energy resulting from mutations in protein complexes. It processes 3D structures by generating residue embeddings and identifying critical interactions between antibody and antigen pairs. By comparing the wild-type and mutated complexes—both provided in PDB format—the model estimates ΔΔG. Its performance is validated via fivefold cross-validation on the SKEMPI 2.0 dataset, covering both single- and multi-point mutations.

RDE-PPI is an unsupervised method that predicts how amino acid mutations affect protein-protein interactions by analyzing changes in side-chain conformations. It uses a flow-based generative model to estimate rotamer distributions and calculates entropy differences between the unbound and bound states to infer binding free energy changes (ΔΔG). The model is trained on structural data from PDB-REDO, where protein chains are clustered and processed into 128-residue patches with introduced noise and masked rotamers, and its ΔΔG predictions are calibrated using the SKEMPI 2.0 database.

### Publicly Available Datasets: SKEMPIv2 and AB-Bind

To the best of our knowledge, the two most widely employed datasets in machine learning tasks related to antibody design/affinity maturation are AB-Bind^9^ and SKEMPI^8^. These datasets collate affinity measurement data points from the PDB that can be associated with particular mutations. The datasets are thus used both as training sets but also to benchmark zero-shot methods that are thought to have implicit correlation with free energy changes. In this study, we used a filtered subset of SKEMPIv2 and AB-Bind, removing all data points other than complete (heavy and light chains) antibody-antigen complexes, followed by a deduplication step.

### Deduplication

Both SKEMPI and AB-Bind contain multiple data points for the same set of mutations. The majority of those duplicates are caused by assaying the same proteins with different methods (e.g. KinExA and ELISA) or at different temperatures (usually 298 K and 310 K). However, even after controlling for those two factors, there are still 52 duplicates in SKEMPI, differing usually by the reference field and affinity value. We noticed that usually one of the duplicates contains detailed measurements (fields: *kon_mut (M^(-1)s^(-1), kon_mut_parsed, kon_wt (M^(-1)s^(-1)), kon_wt_parsed, koff_mut (s^(-1)), koff_mut_parsed, koff_wt (s^(-1)), koff_wt_parsed*) while the other does not. For these cases, we deduplicate by choosing the datapoint with more details. We also deduplicate on *Method* and *Temperature*, because benchmarked models cannot incorporate such information. We choose the row with a more sophisticated Method e.g., KinExA over ELISA, and lower Temperature. In AB-Bind, duplicates are much less frequent, and were handled using a similar logic.

### SKEMPI “train-only” subdataset construction

While all benchmarked ML models were trained on SKEMPIv2, train/test set partitioning strategies varied across the studies underlying model development. This inconsistency prevents fair benchmarking using the full SKEMPIv2 dataset, since the amount of information leakage would not be constant across models.To address this challenge, we constructed a “train-only” subdataset from SKEMPIv2 consisting of the intersection of training data used across all evaluated models. Only the RDE-PPI model provided explicit cross-validation split labels: we selected the training set from fold 0 of the published 3-fold cross-validation, as this fold exhibited the greatest overlap (76%, versus 66% or 58% for folds 1 and 2, respectively) with our filtered SKEMPIv2 Ab-Ag subdataset.).

For DSMBind, the authors reported using the entire SKEMPIv2 dataset for training, excluding the structures already present in SAbDab that were used for testing. We intersected DSMBind’s SKEMPIv2 train subset with our Ab-Ag SKEMPIv2 subset cutting our version to 79% of original size. Finally, for Binding-DDG-predictor, the train-test split procedure was ambiguous: based on available documentation, we inferred that SKEMPIv2 was used for training and SKEMPIv1 for testing, implying that 100% of our SKEMPIv2 Ab-Ag subdataset was included in the training process.

By intersecting the RDE-PPI fold 0 training set (76% of our SKEMPIv2 Ab-Ag subdataset) with the DSMBind training set (79% of our subdataset), we obtained a common set of 336 data points. This intersection represents approximately 59% of our full filtered SKEMPI (Ab-Ag) dataset and serves as the SKEMPI train-only benchmark subdataset for subsequent analyses.

Overall, by constructing the “train-only” subdataset, we enabled comparing ΔΔG prediction models using a set of benchmarks with well-defined train-test leakage, which is often missing or not explicitly controlled for in much of the literature: fully-leaking (SKEMPI “train-only”, a subset of “full”), variable-leaking (SKEMPI “full”, where full means the entire Ab-Ag interaction subset of SKEMPIv2, as described in Methods) ), and non-leaking (AbDesign) datasets.

### Availability

AbDesign DB is available to non-commercial entities for non-commercial use from the following link: naturalantibody.com/ab-design/.

## Results

### AbDesign Curation

The database was constructed by selecting 14 antibodies targeting seven antigens‒two distinct antibodies per antigen (Table 1). Altogether, we did not specifically focus on therapeutic antibodies or potential targets. Some of the targets were of therapeutic interest, such as VEGF or PD-1, but some were simply a reflection of frequently crystallized antigens in the PDB - e.g., lysozyme.

We enumerated atomic contacts between CDR-H3 and antigen residues as those with heavy atoms less than 4.5Å apart. CDR-H3 positions within this distance were then chosen for point mutations (Figure 2). We strived to maximize the number of single mutants, at the same time knowing that the number had to be constrained to 96 antibody variants per plate per target. For this reason, mutations were first automatically distributed to maximize the number of mutations per position, whilst not favoring any position over another. Each set of mutations was then manually checked to identify mutations that would appear clearly non-informative. For instance, the introduction of cysteines was avoided, and in certain cases, proline or alanine mutations were not introduced because of structural variability or overrepresentation in other datasets. This procedure resulted in 658 mutations with an average of 47 mutations per antibody. The full statistics of the dataset are given in Table 1.

**Figure 2.**
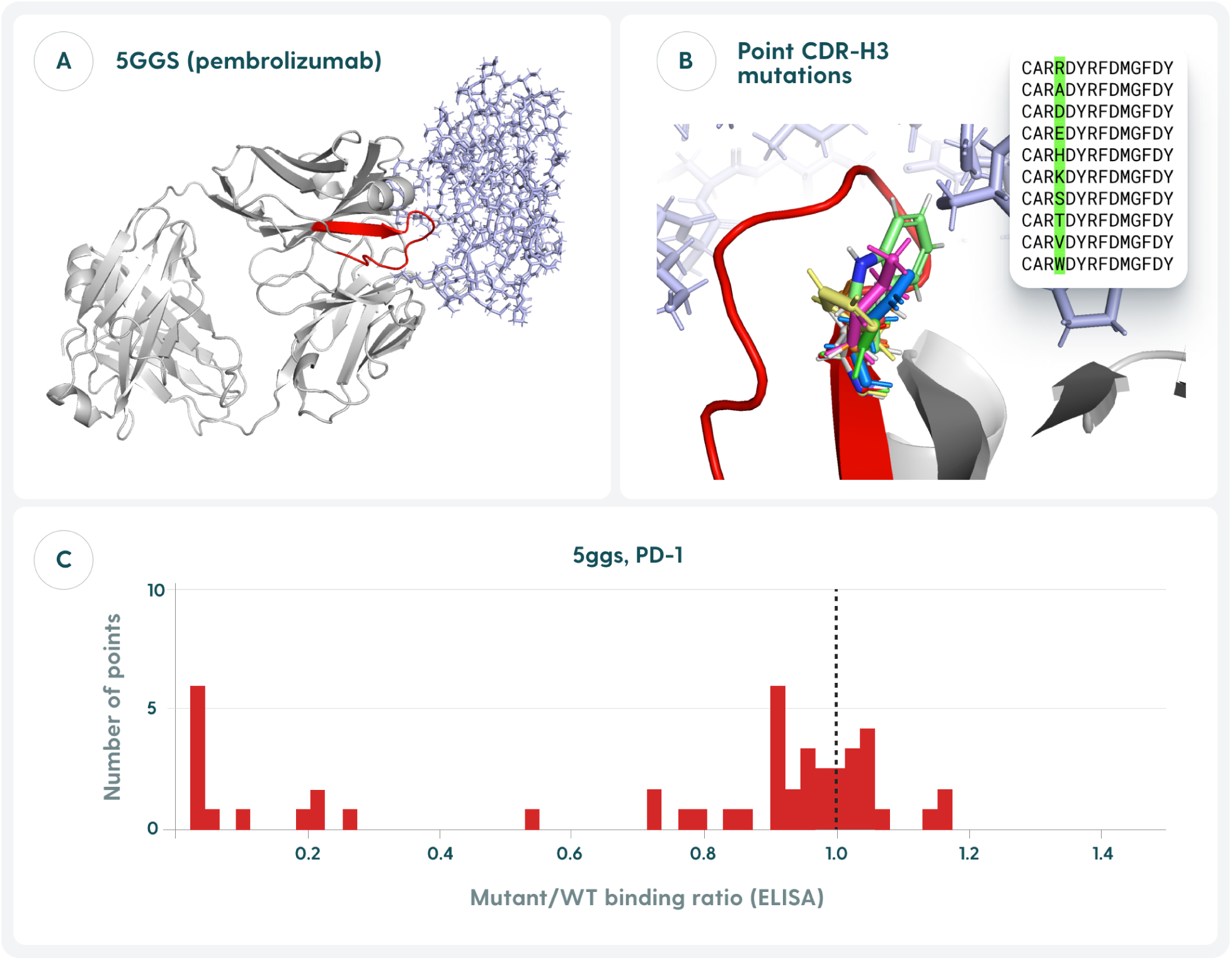
The mutational strategy of AbDesign. **A.** We selected complexes from the Protein Data Bank such that at least two distinct antibodies were available against a single target. **B.** The contact residues (<4.5Å of CDR-H3 heavy atom to an antigen heavy atom) were subjected to point mutations. **C.** The variant binding affinity to the target was measured using ELISA, having the wild-type readout on the same plate. The ratio between the readout of the variant to that of the wild-type (black vertical line) indicates whether binding affinity was worse (<1) or not (≥1) than the wild-type reference. The use of natural ratios as opposed to log ratios is motivated by our intention to highlight loss of binding.

We envisage the usage of the database in affinity maturation experiments where the introduced mutations must not destroy the wild-type binding. Though there is no guarantee that the mutant maintains the same shape as the parent, it is a feasible assumption. Therefore, we modeled each structure using ABodyBuilder2^16^ and performed the minimization of it using OpenMM. We tagged each file with the mutation that was applied. Such structural data can thus be linked to sequence and ELISA ratios, making the dataset better suited for machine learning applications.

### Contrast between SKEMPI, AB-Bind, and AbDesign

We contrast publicly available SKEMPIv2 and AB-Bind datasets with ours on two levels: total number of data points and the distribution of mutations.

Table 4 offers the raw number of data points in AbDesign, SKEMPI, and AB-Bind. SKEMPI is the most plentiful resource, as it also includes non-antibody data points. Both resources include mutations on the antigen as well, so for comparison with ours, we had to constrain these to only the structures where the antibody was modified. Furthermore, both datasets contain a number of multiple mutants, whereas we focus solely on point mutants. To better understand the relationship between the datasets, we also assessed the overlap at both the PDB and mutation levels. A Venn diagram of PDB IDs (Supplementary Figure 1) reveals that SKEMPI and AB-Bind share 11 structures, while AbDesign is nearly non-overlapping, with only one structure shared with SKEMPI. We further examined the specific mutations shared between SKEMPI and AB-Bind and found that, although some overlap exists, it is relatively limited, with 57 shared mutants (Supplementary Figure 2). These observations highlight that AbDesign provides a structurally and mutationally independent test set, which is particularly valuable for benchmarking machine learning models trained on SKEMPI, as it enables an unbiased evaluation of generalizability.

**Table 4.** Contrast of raw data points between SKEMPI, AB-Bind, and AbDesign. SKEMPI and AB-Bind collated mutations might be on the antigen side. SKEMPI had non-antibody data points as well. The antibody-only data points reflect the filtered versions of both SKEMPI and AB-Bind.

|  |  | Database |  |  |  |
| --- | --- | --- | --- | --- | --- |
|  |  | SKEMPI<br>(train only) | SKEMPI<br>(full) | AB-Bind | AbDesign |
| All data points | PDBs | - | 345 | 32 | 14 |
|  | Variants | - | 12,324 | 1,193 | 658 |
| Antibody-only data points | PDBs | 19 | 34 | 13 | 14 |
|  | Variants | 261 point mutants<br>(336 all) | 377 point mutants<br>(571 all) | 268 point mutants<br>(351 all) | 658 point mutants |

The second major point of contrast beyond raw numbers is the types of mutations that we targeted and that are introduced. In Figure 3, we show what residues are used to point mutate (top) and what amino acids are used as replacements (bottom). The apparent bias in tyrosines (Y) that are chosen for mutations in all three datasets simply reflects the overrepresentation of this particular residue in binding, which has been widely documented ^22,23^. The mutational bias for alanine in AB-Bind and SKEMPI can be easily attributed to the multiple alanine scanning experiments that the studies captured. By contrast, the mutations that we introduced are more evenly distributed. Though we note a particular undersampling of L, M, N, P, and Q due to the sampling procedure that favored all other amino acids before proceeding to these.

**Figure 3.**
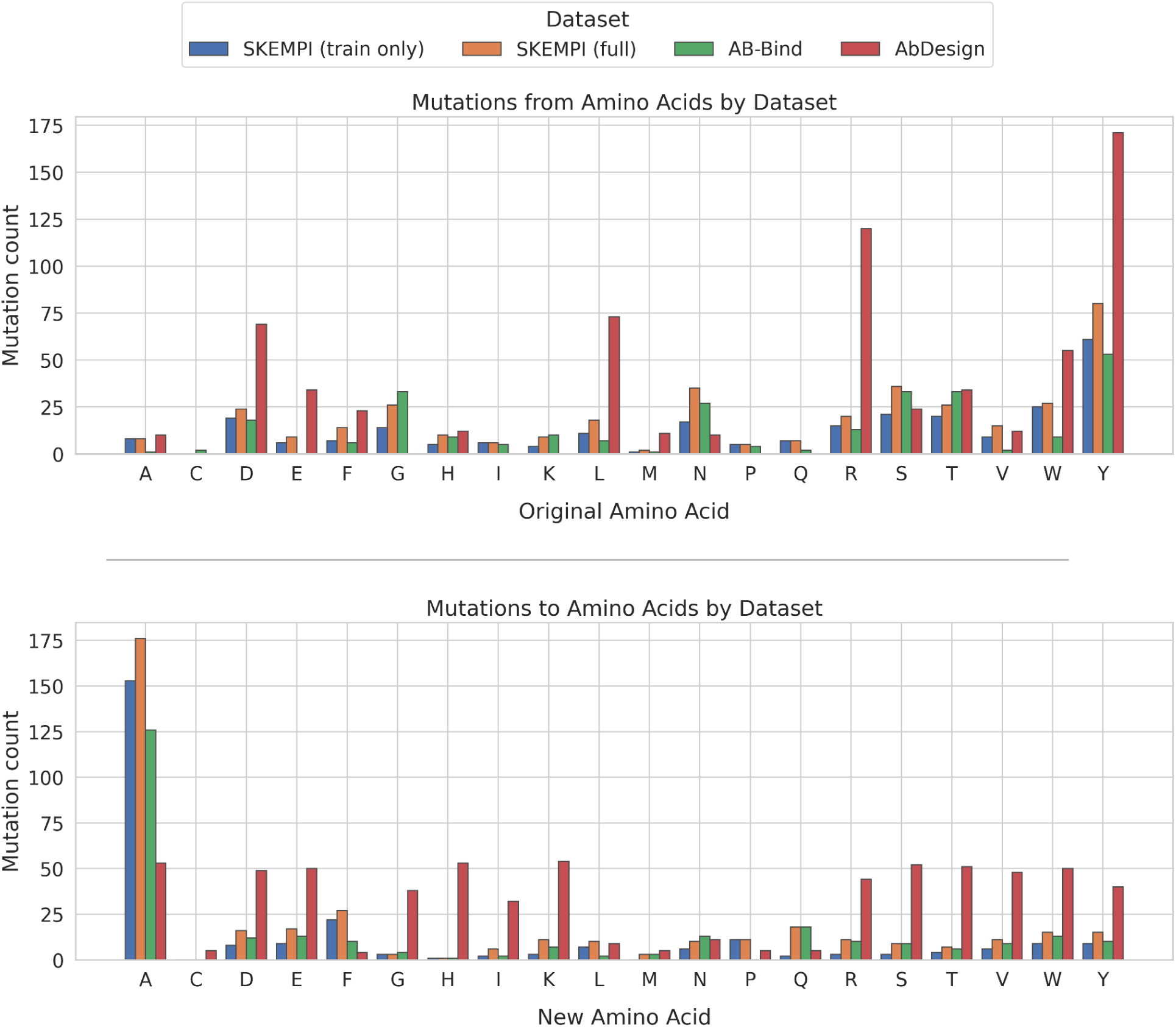
Contrast between point mutation types across datasets. The data presented is for the Ab-Ag subsets of the SKEMPIv2 and AB-Bind datasets (Table 4). For SKEMPI, we also show a “train only” subdataset (see Methods). For point mutants in each dataset, we noted how often an amino acid was being picked for mutation (top) and how often an amino acid was being selected as replacement (bottom).

When considering only the raw number of point mutants, our dataset is currently the most plentiful, though still small, given the scales needed to employ such resources for training ^24^. Furthermore, our dataset offers novel binding data points generated under single experimental conditions rather than points collated from different laboratories. The chief limitation of our dataset is that it only offers ELISA readouts rather than more precise affinity measurements such as SPR^25^.

### Most Complexes are Resilient to Point Mutations

For each antibody complex, we had an ELISA readout for the wild-type antibody (WT) and the mutant at a particular concentration. The ratio between these two values was employed as a proxy to gauge whether binding was destroyed with respect to the WT or not - we term it ‘binding retainment’. Mutations with binding retainment of 1 have identical ELISA profiles to the wild-type. Those approaching 0 have lost binding with respect to the wild-type.

In Figure 4, we plotted the histograms of the binding retainment for each of the 14 PDBs. We note that in most cases, there is no clear separation between binders and non-binders, with the binding retainment values either being centered around wild-type or closer to 0, but with some values in between. One notes a wide spread of values, indicating that point mutations have wide-ranging effects rather than simply maintaining or knocking off binding altogether. However, in the case of 1BVK, all mutations can be classified as detrimental, whereas in the case of 6RP8 as neutral.

**Figure 4.**
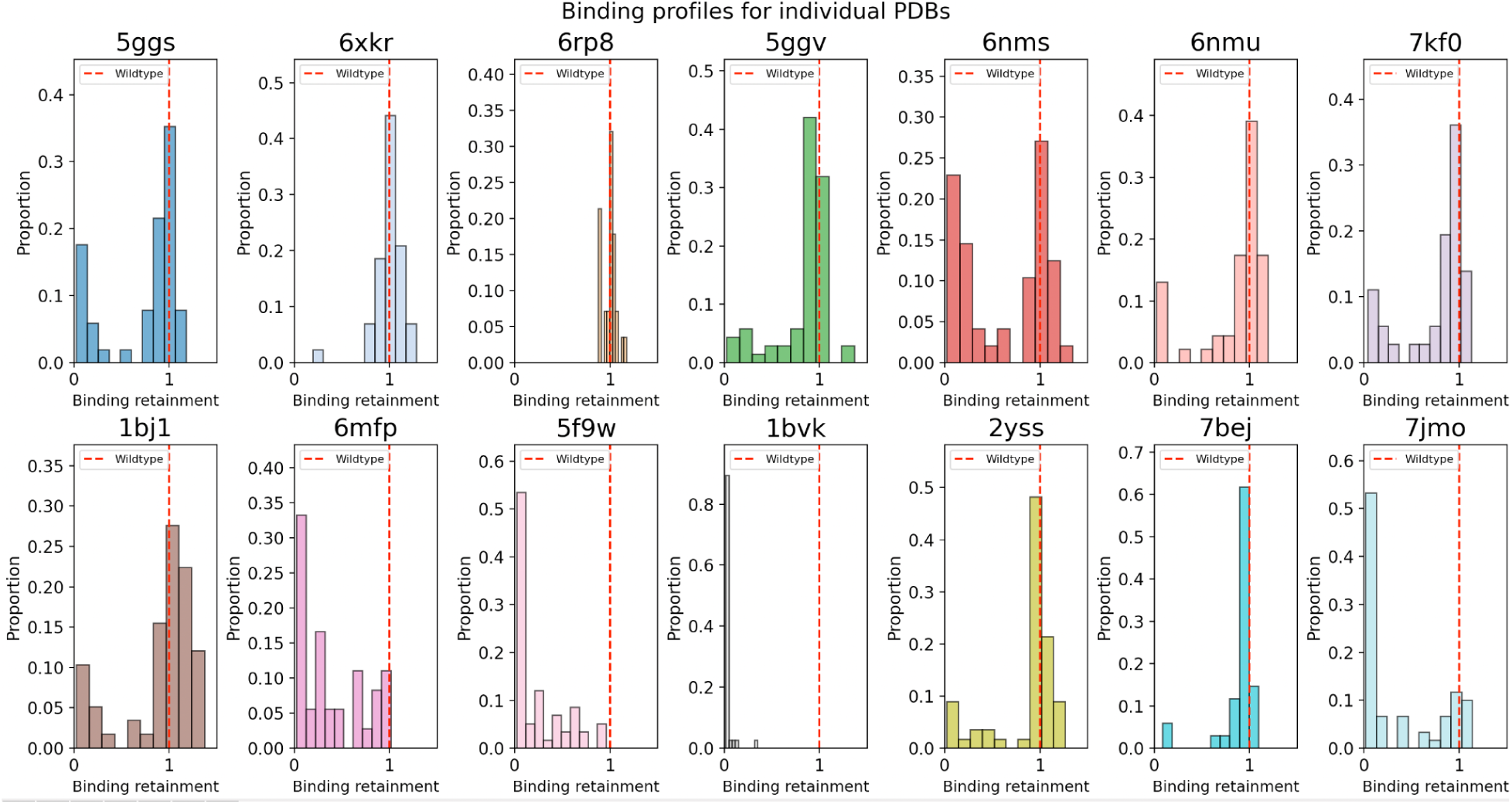
Binding profiles for the point mutants for each of the 14 PDBs of the AbDesign dataset. We selected seven antigens, with two PDBs per antigen, with distinct antibodies. Each antibody CDR-H3 was mutated, and we noted the ratio of its ELISA binding to the wild-type (vertical black line = 1). Ratios closer to 1 indicate a similar ELISA profile as wild-type, whereas values approaching 0 can be indicative of reduced binding. While log-transformed ratios are commonly used to ensure symmetry in up- and down-regulation, here we report raw ratios to preserve interpretability in the context of binding loss. Since our primary focus is on decreases in [binding/activity/interaction], retaining ratios in their original scale (where values <1 denote loss) provides a more direct and intuitive representation of the effect magnitude.

To build a better understanding of the number of mutations that are detrimental/favorable to binding in such point-mutagenesis exercise, we plotted the number of residues that are classified as ‘binding’ for a number of thresholds of the binding retainment value in Figure 5. In most cases, the binding sites are quite resilient to modifications. Taking a high value of 1 (wild-type) as a threshold and performing random mutations results in 26% mutations on average retaining binding. The ratio jumps to 51% when we consider binding retainment at a level of 0.9, which might be slightly weaker than wild-type, but still within the binding range. In computational terms, it means that performing random point mutagenesis to a wide range of amino acids (not only alanines) results in a seemingly high random baseline accuracy of around 50%.

**Figure 5.**
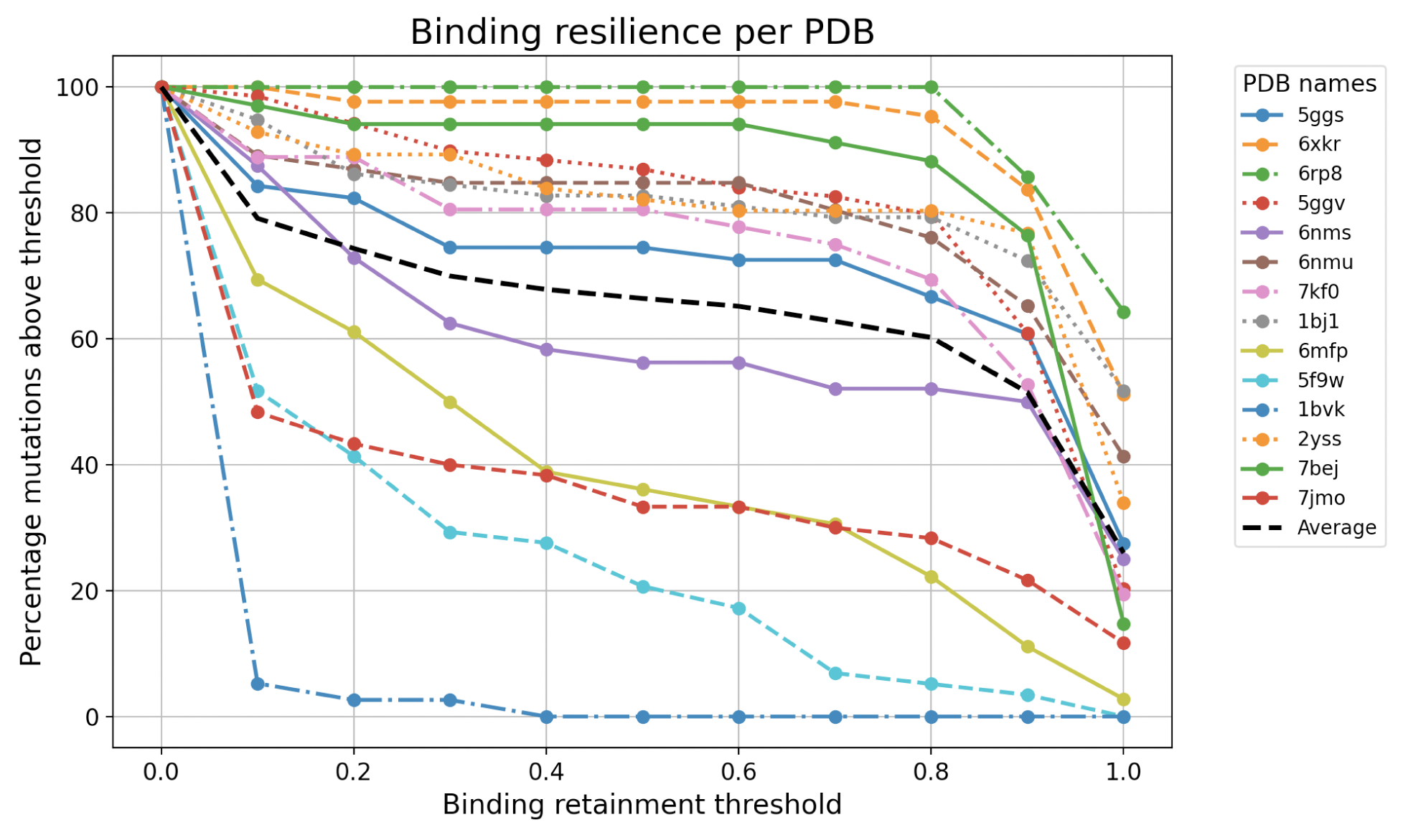
Binding resilience of point mutants in AbDesign. For each PDB, we plotted the percentage of mutants with ELISA ratio above the given threshold. The wild-type for each PDB has a value of 1, and values below it can be interpreted as weaker binding. Values approaching zero are thought to be non-binding.

### Benchmarking Machine Learning-based Energy Models

By contributing the AbDesign DB resource, we are in a position to offer a blind hold-out set for a number of machine learning models that predict the effects of binding. Indeed, algorithms are all generally trained on various folds of SKEMPI/AB-Bind. We benchmarked three models that predict the binding strength of a complex and its mutants: RDE-PPI, Binding-DDG-predictor, and DSMbind. The choice of the methods was motivated by performance, code availability and license permissibility. In their original scientific publications, all three methods showed acceptable correlation between predictions and experimental measurements of binding energy of antibody variants as benchmarked using SKEMPI and/or AB-Bind: however, noting that these methods were also trained on these datasets.

We tested the ML algorithms on our curated Ab-Ag subsets of SKEMPI (full and “train only”, see Methods) and AB-Bind (Table 4) to ensure that we are reproducing the protocols correctly, with results given in Supplementary Table 1 (Figure 6) & Supplementary Table 2 (Figure 7). We report the correlations with the affinity data points in a per-PDB averaged data (Supplementary Table 1) and with all the data points taken together (Supplementary Table 2). Note, the SKEMPI and AB-Bind datasets include both single- and multi-point mutations, whereas the AbDesign dataset consists of only single-point mutations.

**Figure 6.**
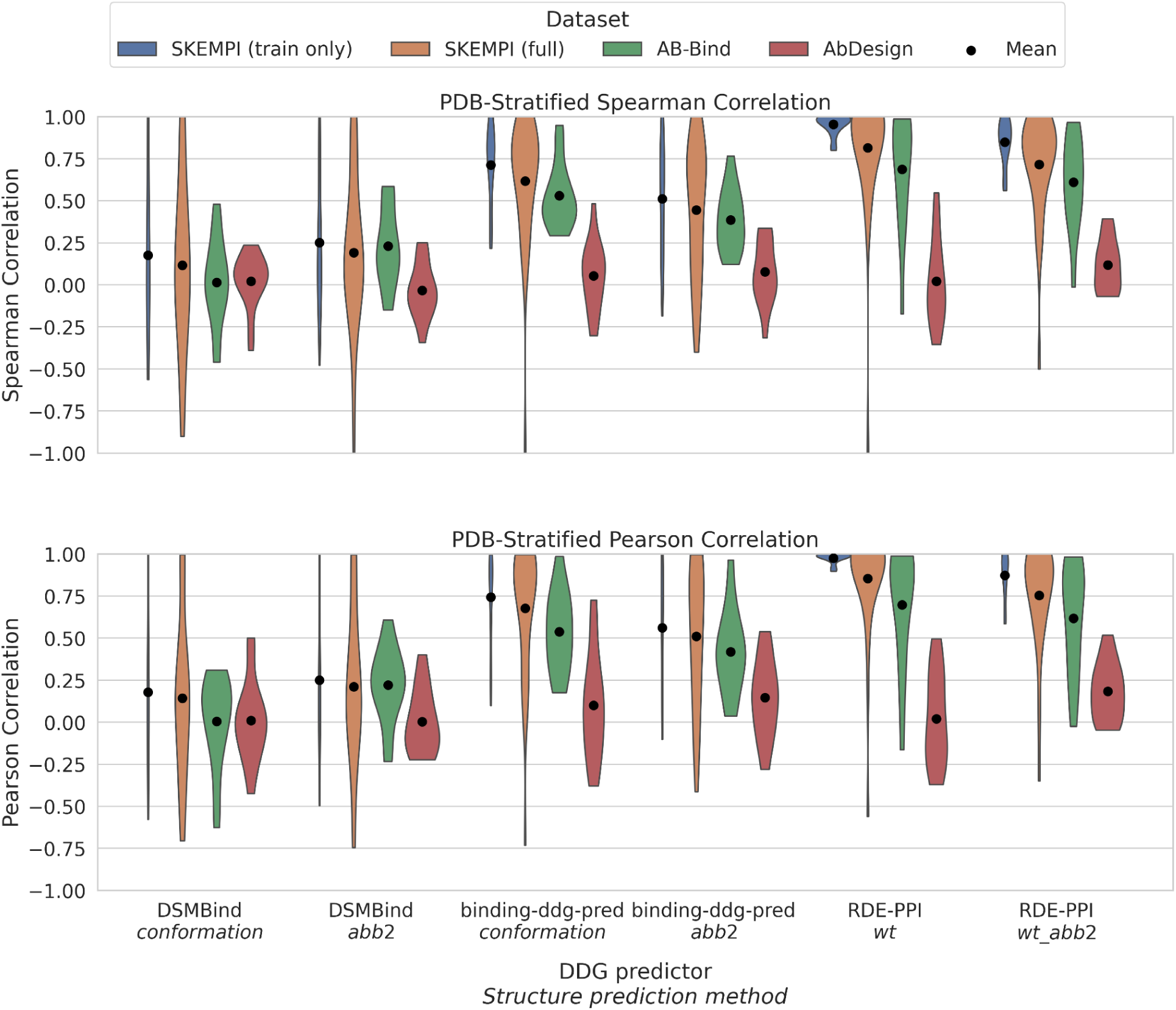
PDB-stratified Pearson and Spearman correlation distributions across datasets. This figure presents violin plots of the Pearson (top) and Spearman (bottom) correlation distributions between model scores and experimental values for different dataset-structure prediction method combinations. Each distribution represents per-PDB correlation values for a specific structure prediction method and dataset. The models analyzed include DSMbind, Binding-DDG-predictor, and RDE-PPI, while the datasets include SKEMPI (train only), SKEMPI (full), AB-Bind, and AbDesign. The train-only variant of SKEMPI is an intersection of the train datasets of all models. The black dots indicate the mean correlation value for each group. For the AbDesign dataset, the sign of the correlation values has been reversed to match the scale direction of the SKEMPI and AB-Bind datasets.

**Figure 7.**
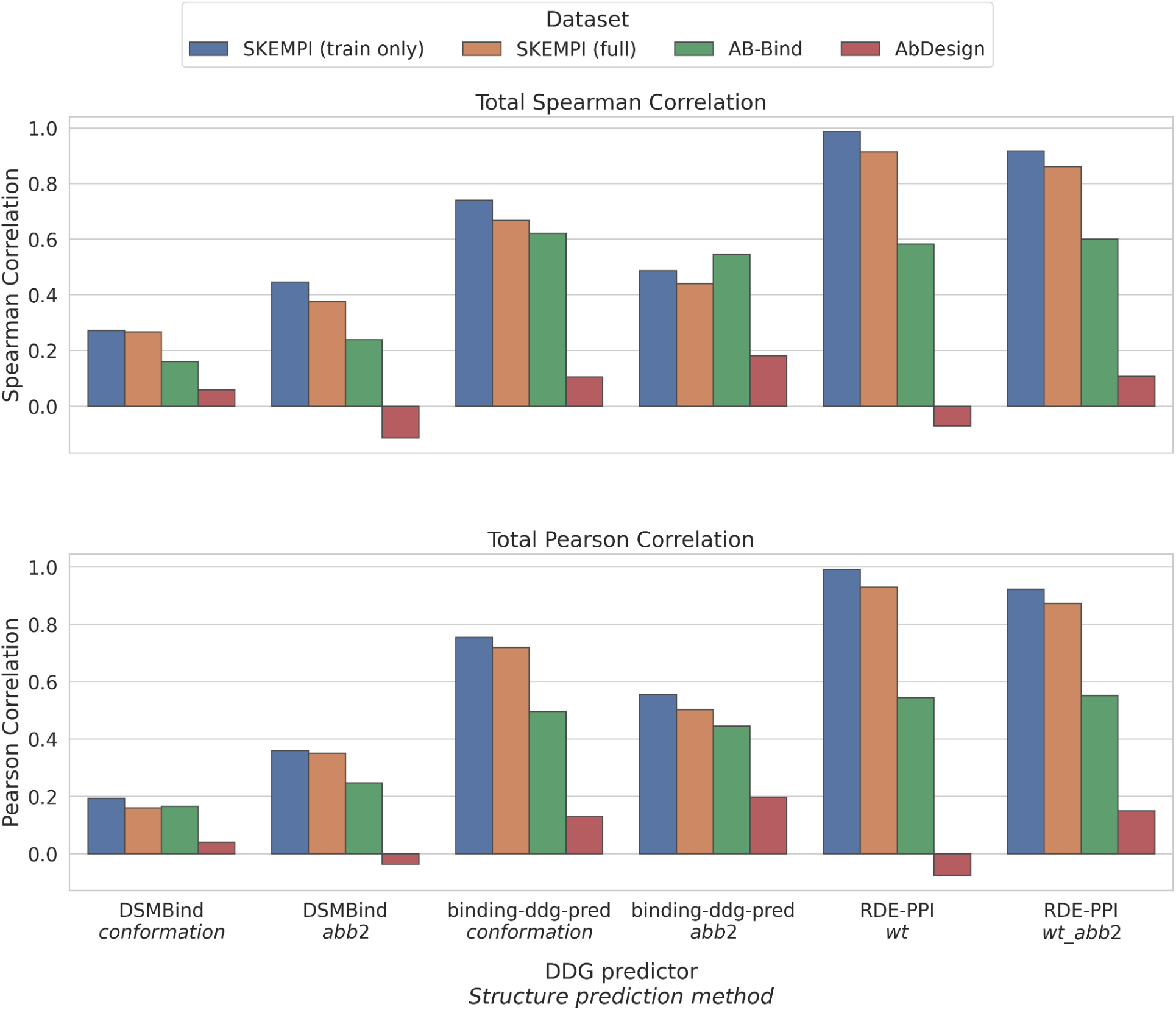
Total Correlations, without PDB stratification. Bar plots of the Pearson (top) and Spearman (bottom) correlation values between model scores and experimental values for different dataset-modeling method combinations. Each bar represents the total correlation value per modeling method and dataset group. The models analyzed include DSMbind, Binding-DDG-predictor, and RDE-PPI, while the datasets include SKEMPI (train only), SKEMPI (full), AB-Bind, and AbDesign. The train-only variant of SKEMPI is an intersection of the train datasets of all models. For the AbDesign dataset, the sign of the correlation values has been reversed to match the scale direction of the SKEMPI and AB-Bind datasets. In AbDesign, lower ELISA ratio values indicate loss of binding, while values greater than 1 indicate gain, opposite to the ΔΔG interpretation used in the other datasets.

The models return ΔΔG prediction - a change in Gibbs free energy (ΔG) upon mutation (wild-type vs mutant) and binding (unbound vs bound). We define ΔG as:

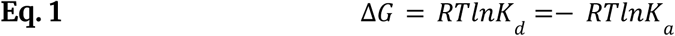

Where:

ΔG = Gibbs free energy change (kcal/mol or kJ/mol)

R = Universal gas constant (1.987 cal/(mol·K) or 8.314 J/(mol·K))

T = Absolute temperature (Kelvin, typically 298 K for room temperature) K_d_ = Dissociation constant (Molar, M)

K_a_ = Association constant (Molar, M)

ΔΔG represents the difference in binding free energy before and after a mutation:

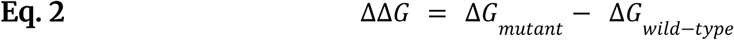

Where:

- **If ΔΔG > 0**, the mutation decreases binding affinity (weaker binding).
- **If ΔΔG < 0**, the mutation increases binding affinity (stronger binding).
- **If ΔΔG = 0**, the mutation does not affect binding.

If the predicted ΔΔG is positive, it should correspond to the cases where binding was reduced. Conversely, negative prediction should correlate with preservation / strengthening of binding.

Binding energy estimations are dependent on structural predictions, as mutants are not guaranteed to assume the conformation of the wild-type antibody. For this reason, we tested two structure prediction methods: homology-based side chain remodeling and machine-learning ABB2, which we named ‘conformation’ and ‘abb2’, respectively.

Conformational replacement involves picking the most common side chain based on density, as observed in the PDB, followed by OpenMM refinement. In the case of RDE-PPI, which receives only the WT structure alongside mutation specification, we also used the experimental structures to further contrast its performance with structure prediction methods.

Altogether, the methods perform well on the datasets they were trained on, reflecting that we are using the protocols correctly. Nevertheless, the prediction-vs-measurement correlations on the AbDesign dataset are very poor, with the Spearman’s and Pearson’s correlations in the range of 0.1. The results are consistent if we take individual PDBs into account for energy correlations (Supplementary Table 1 & Figure 6) as well as when we take the entire dataset holistically (Supplementary Table 2 & Figure 7). Altogether, this indicates that the more heterogeneous AbDesign dataset, that is a proper hold-out, is more challenging for methods to generalize to. We also note a wide variation depending on whether we employed the conformation or ABB2-based modeling. It is not surprising, however, indicating the dependence of the ΔΔG prediction methods on accurate structures.

RDE-PPI performed the best and the most consistently across the datasets, especially if we take the performance on the AbDesign dataset into account. This method was trained on three different folds of SKEMPI, and in Figure 8, we show the variation of its predictions. The predictions across the folds are consistent for SKEMPI and AB-Bind, which is not surprising given that the dataset was trained on these. The worst performance is on the experimental structure dataset in AbDesign. By contrast, the results for the folds are consistent for the predicted structures of the antibodies from the same dataset.

**Figure 8.**
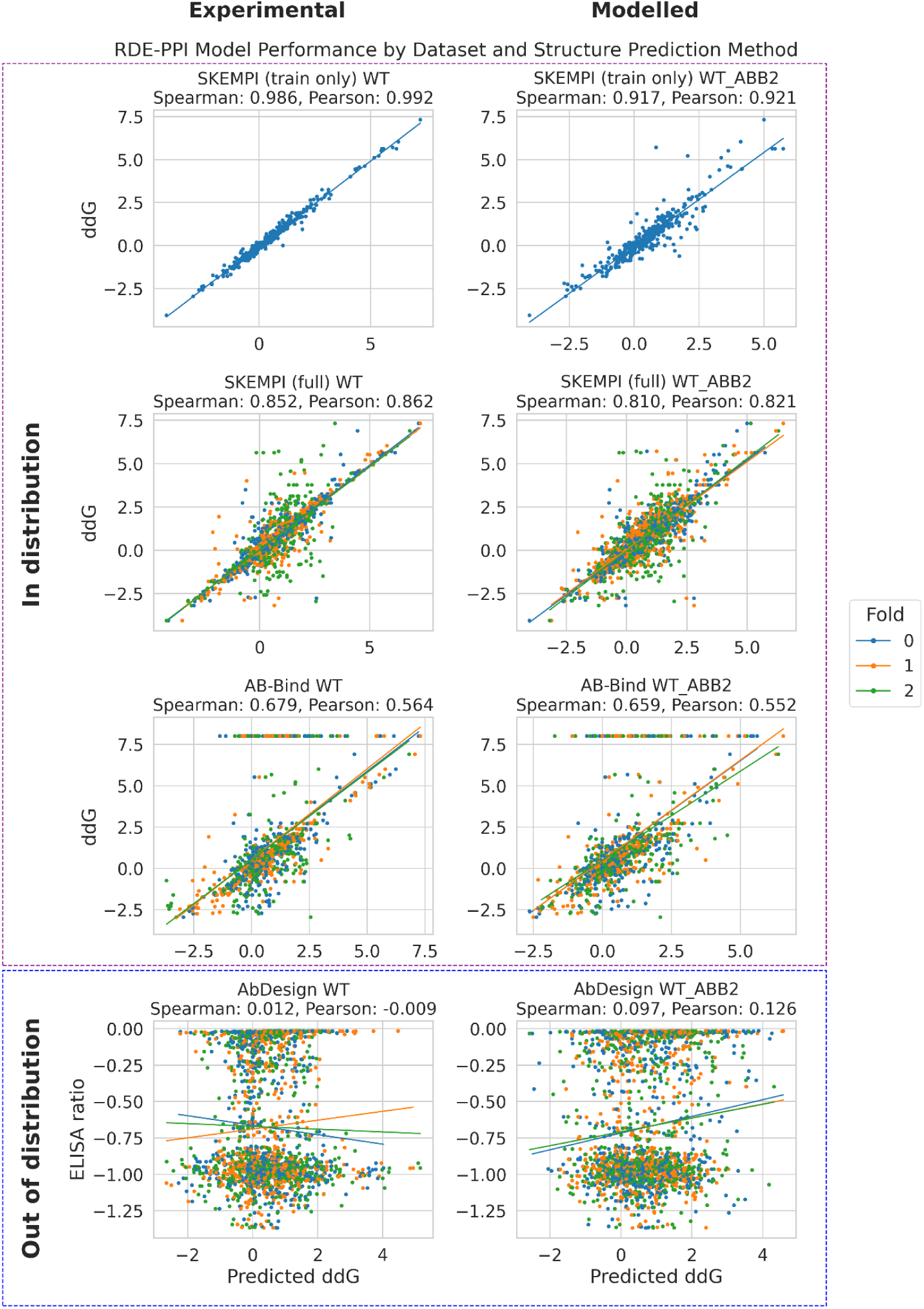
Variation of folds of RDE-PPI on the predictive performance. RDE-PPI was trained using three folds of SKEMPI (0, 1, 2). We show the correlation of predicted versus actual ΔΔG (SKEMPI or AB-Bind) or ELISA binding ratios (AbDesign). The “train only” subset of SKEMPI was extracted using fold 0 of the RDE-PPI dataset, hence the absence of data points from other folds in the first plot row.

We investigated whether stratification by mutation type might reveal some biases of the models/datasets used for training. Figure 9 demonstrates performance for different models/datasets per mutation type (biochemical and size). It is clear from Figure 3A that tyrosine is the most commonly mutated residue, as a result of its known overrepresentation in antibody binding sites. Other residues have much more uneven representation. The most common mutation is alanine, as apparent from Figure 3B. Alanine scanning is a very common technique, and the data reflects it. Given the small numbers we are faced with, we find that stratifying correlations by individual mutants might not lead to clear statistical conclusions, even less so for the mutant pairs. The only clear signal we tested was indeed the alanine mutation. In Supplementary Table 3 (Figure 10 and Figure 11), we contrasted the predictive power of a model mutating to alanine versus any other amino acid. We find that the models do much better on predicting alanine mutants than other modifications, which highlights that models do not generalize well beyond the data they were given at training time.

**Figure 9.**
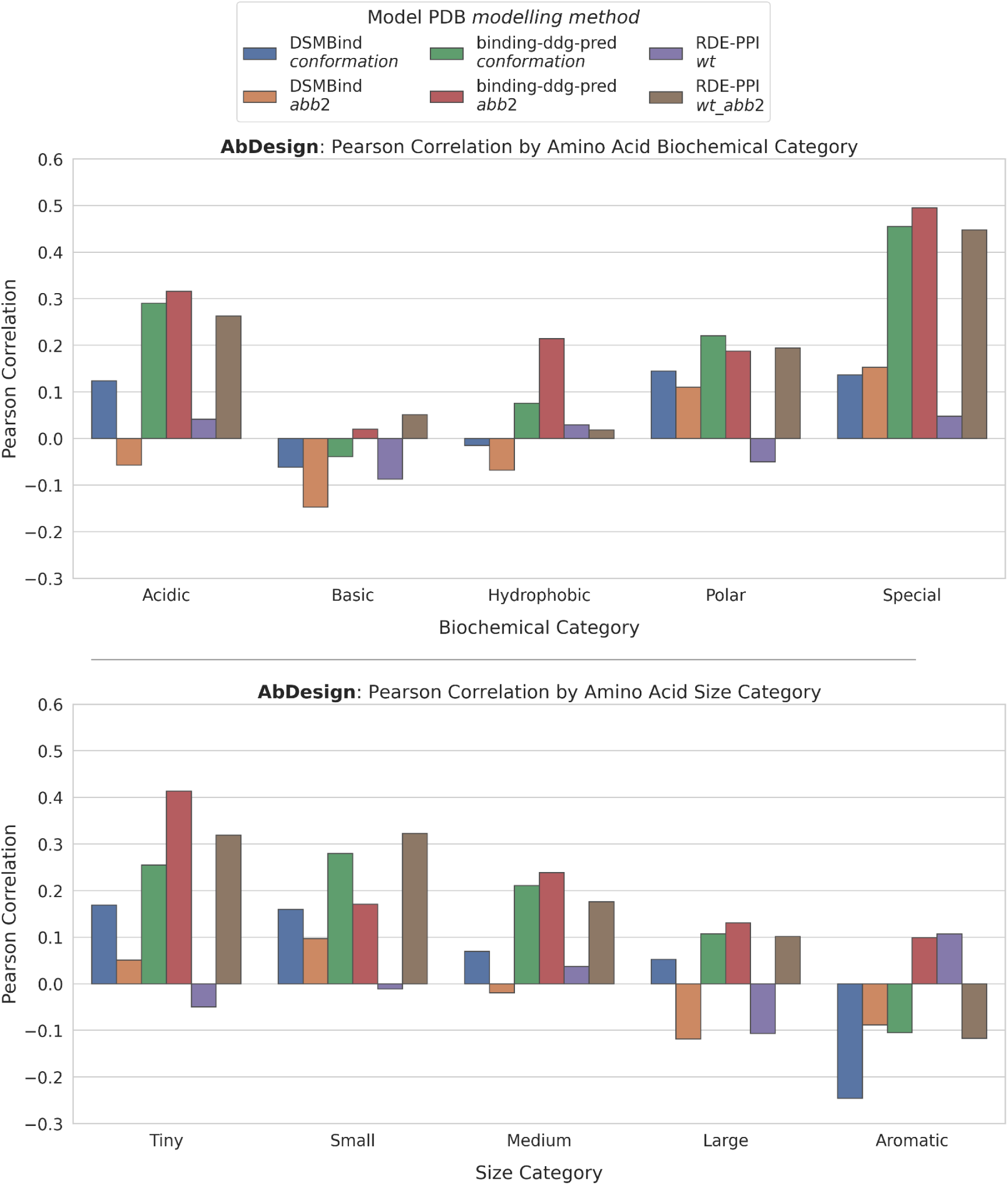
Benchmarking models on AbDesign grouped by mutated amino acid properties. This figure presents Pearson correlation values between model-predicted and experimentally measured binding effects, restricted to the AbDesign dataset and stratified by properties of the mutated (new) amino acid. The top panel shows correlations grouped by biochemical properties of the substituted residue (Acidic, Basic, Hydrophobic, Polar, Special), while the bottom panel shows correlations grouped by size (Tiny, Small, Medium, Large, Aromatic). Bars represent total correlation values per model and grouping. The modeling methods compared include DSMbind, Binding-DDG-predictor, and RDE-PPI, evaluated across two model PDB conformations (`conformation` and `abb2;` or `wt` and `wt_abb2`). See Supplementary Table 4 for the amino acid groupings used in each category.

**Figure 10.**
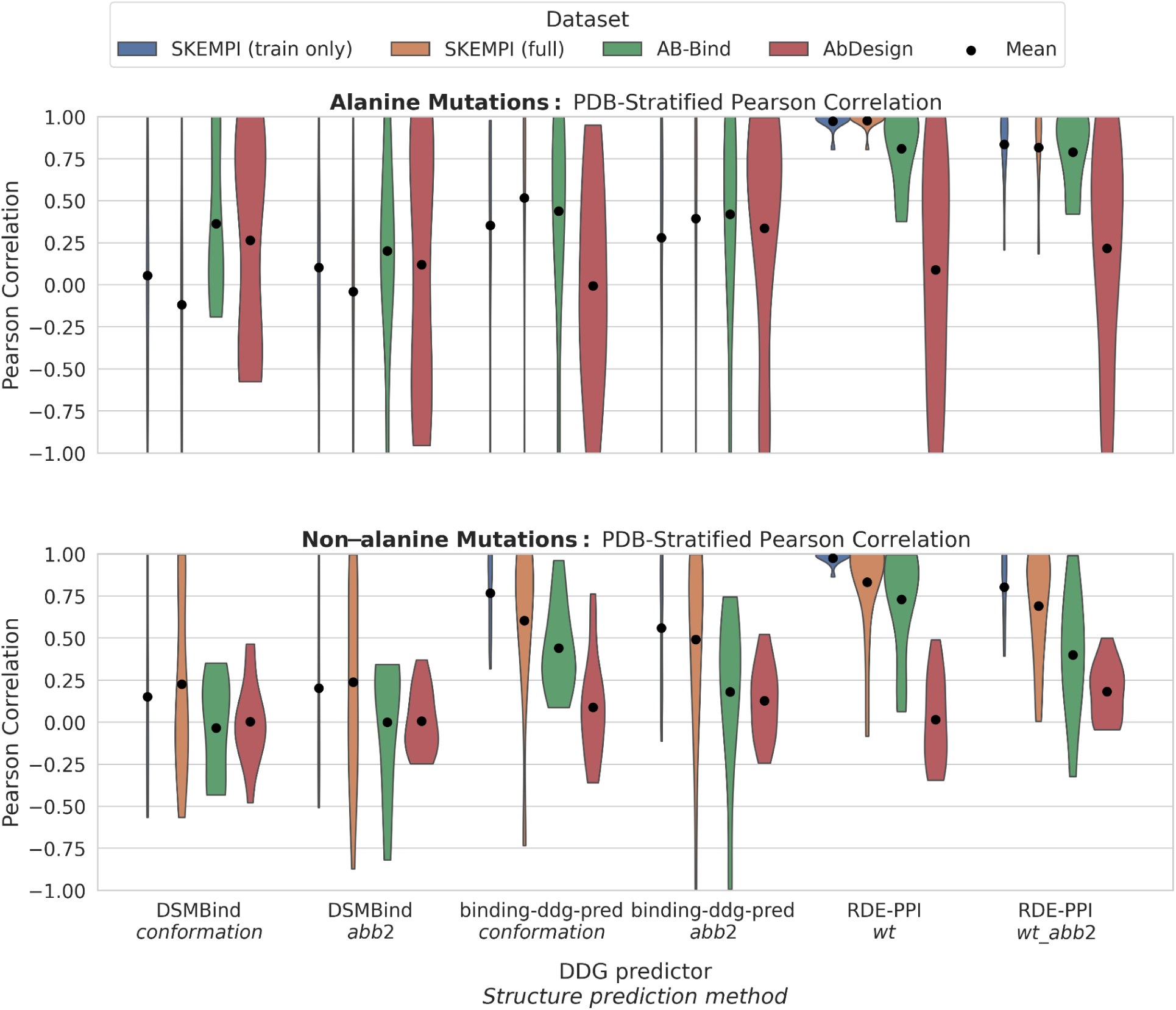
PDB-specific Pearson correlation distributions for mutations to Alanine vs. non-Alanine mutations. This figure shows violin plots of the Pearson correlation distributions between model scores and experimental values for different dataset-modeling method combinations. The top panel displays correlations for mutations to Alanine, while the bottom panel covers all other mutation types. Each distribution represents per-PDB correlation values for a specific modeling method and dataset. The models analyzed include DSMbind, Binding-DDG-predictor, and RDE-PPI, while the datasets include SKEMPI (train only), SKEMPI (full), AB-Bind, and AbDesign. The black dots indicate the mean correlation value for each group. For the AbDesign dataset, the sign of the correlation values has been reversed to match the scale direction of the SKEMPI and AB-Bind datasets. In AbDesign, lower ELISA ratio values indicate loss of binding, while values greater than 1 indicate gain, opposite to the ΔΔG interpretation used in the other datasets.

**Figure 11.**
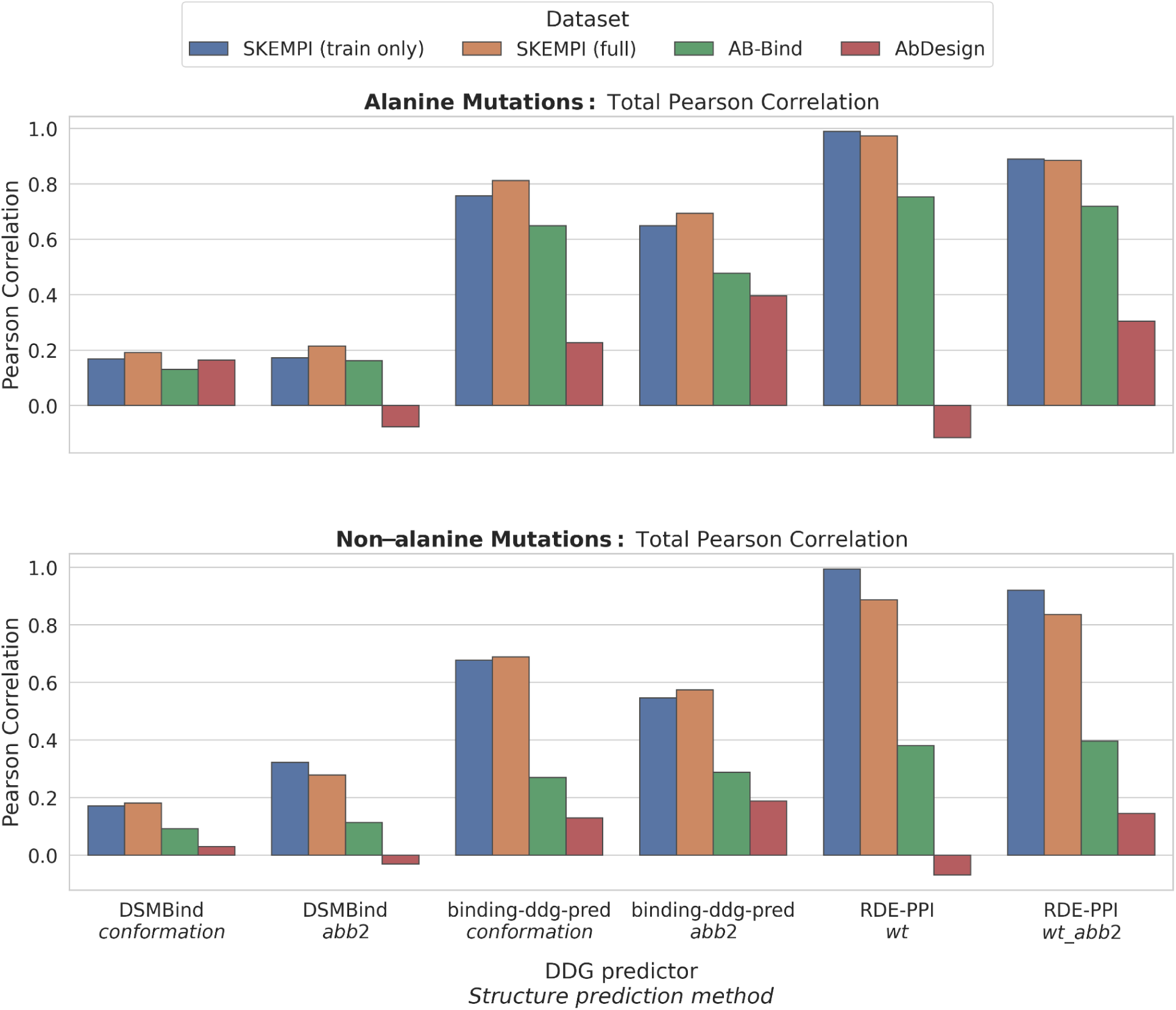
Total Pearson correlations given Alanine mutations. Bar plots showing the Pearson correlation between model scores and experimental values across different dataset–modeling method combinations, separated by mutation type. The top panel presents correlations for mutations to Alanine, while the bottom panel includes all other substitutions. Each bar represents the total Pearson correlation for a given modeling method and dataset. The models evaluated are DSMbind, Binding-DDG-predictor, and RDE-PPI, and the datasets include SKEMPI (train only), SKEMPI (full), AB-Bind, and AbDesign. For the AbDesign dataset, the sign of the correlation values has been reversed to match the interpretation of SKEMPI and AB-Bind values.

### Good Correlations between FoldX Predictions and AbDesign Measurements

We chose FoldX as the prominent empirical method to compare the machine learning methods against^26^. Since FoldX was commercially unavailable at the time of conducting this work, we leveraged a pre-computed dataset of FoldX values^18^. Here, we identified nine out of our fourteen PDBs in AbDesign with the mutations we required.

Plots of the FoldX scores against our binding retainment are given in Figure 12. We note consistency in predictions on our dataset, unlike DSMBind, Binding-DDG-predictor, and RDE-PPI. Comparing Pearson’s correlations (best for each method) in Supplementary Table 5 shows the superiority of FoldX on our dataset. For completeness of comparison in Supplementary Figure 3 we demonstrate correlations to FoldX for alanine vs non-alanine mutations This stands to show that some non-ML methods that explicitly take energy calculations into account can generalize better to novel datasets and be useful for design applications.

**Figure 12.**
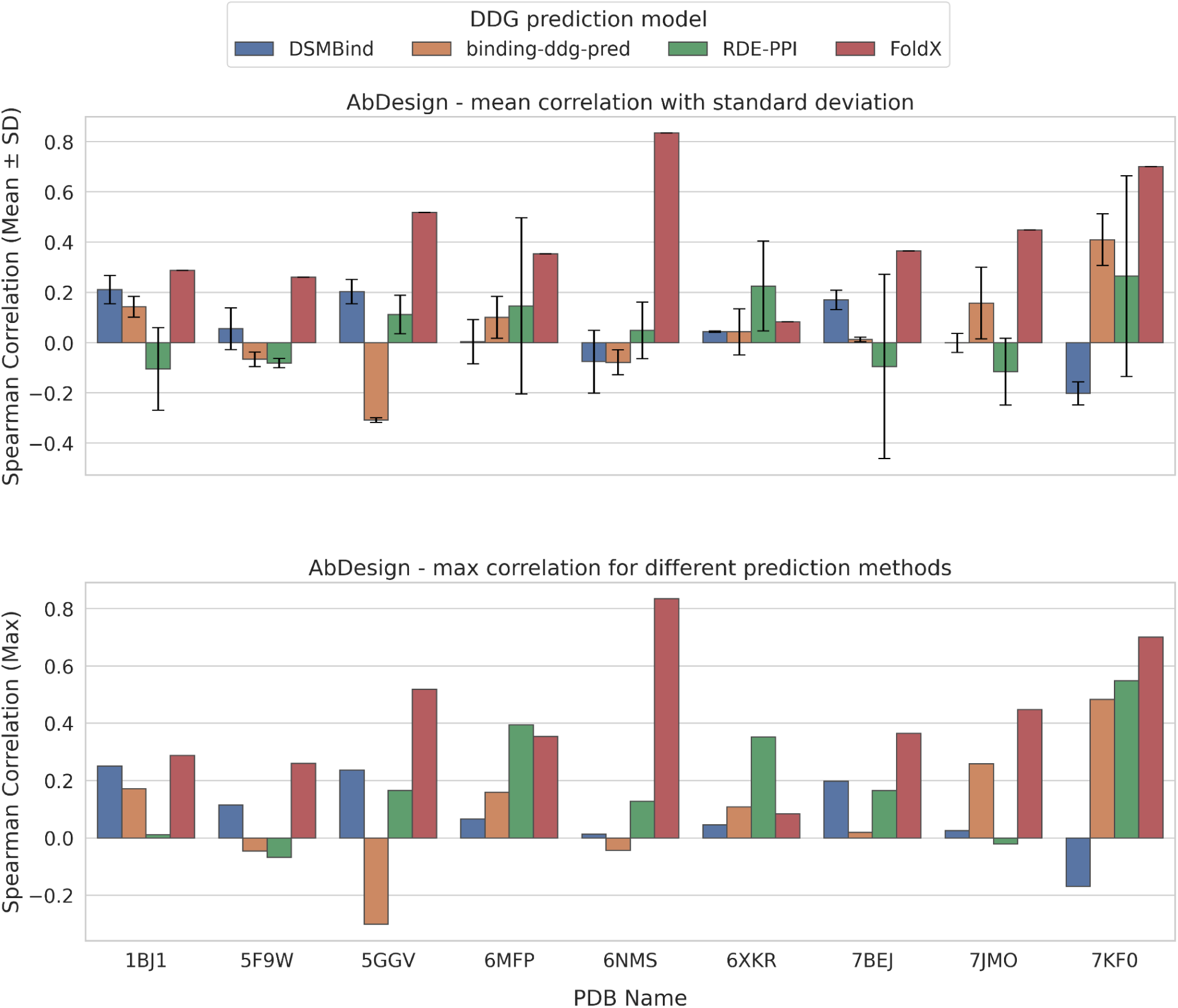
Scores on the reduced FoldX dataset. For 9 out of 14 PDBs in our AbDesign dataset, we could source FoldX predictions from published datasets. For each individual PDB, we show the Spearman’s correlation coefficient between the model score and the binding retainment score. Mean (top panel) and max (bottom panel) correlations across structure prediction approaches are shown for each ML model. As FoldX uses its own structure prediction approach, leading to a single prediction, the mean and max correlations are identical.

## Discussion

Designing antibodies wholly *in silico* has made steady progress^27^, yet its ultimate success remains limited by the availability of structurally annotated, experimentally uniform affinity data ^24^. AbDesign addresses this deficit by providing 658 single-point CDR-H3 mutants—each assayed under an identical ELISA protocol and linked to either high-resolution crystal structures or consistently refined models—thereby more than doubling the number of structurally defined antibody mutants available for benchmarking.

When evaluated on their native datasets (SKEMPI, AB-Bind), modern machine-learning predictors (DSMBind, Binding-DDG-predictor, RDE-PPI) achieve moderate correlations with experimental affinity changes (Spearman’s ρ ≈ 0.4–0.7). However, on AbDesign—comprising entirely non-overlapping PDB complexes—their performance falls to near zero (ρ ≈ 0.0–0.1). This decline highlights two intrinsic limitations. First, existing repositories are dominated by particular mutation types, especially alanine-scanning, which induces training biases. Second, many algorithms exhibit pronounced sensitivity to subtle inaccuracies in mutant side-chain structure prediction, whether generated by deep-learning tools such as ABodyBuilder2 or by classical rotamer replacement followed by energy minimization.

By contrast, the empirical, force-field–based program FoldX maintains reasonable predictive power on AbDesign (Pearson’s R ≈ 0.3–0.5). This resilience recalls the seminal work of Lippow et al. (2007)^28^, who first employed the CHARMM force field to guide point-mutational design and reported reliable identification of binding-preserving substitutions. That simple, physics-driven energy terms—van der Waals interactions, electrostatics, and solvation—continue to generalize to novel antibody–antigen pairs suggests that first-principles approaches remain indispensable, even as machine learning methods advance.

Our results thus expose a gap between “first-principles” and purely data-driven methods when confronted with genuinely new sequence–structure contexts. Machine-learning models perform best within the confines of their training distributions, but can falter on unseen antibodies, which often feature unusual CDR loop lengths and unique somatic-mutation patterns. In contrast, physics-based tools rely on universal chemical principles that transcend individual datasets.

Emerging hybrid frameworks offer a promising route to bridge this divide. For example, METL^29^ integrates empirical energy-term features directly into a transformer architecture, allowing the model to learn both statistical patterns from large structural corpora and fundamental energetic contributions. Such hybrid approaches have already demonstrated improved generalization across diverse protein–protein interfaces and may yield greater robustness on hold-out sets like AbDesign.

Beyond benchmarking, AbDesign has practical implications for antibody engineering. In affinity-maturation campaigns, rapid triaging of candidate mutations is essential. Our findings suggest that force-field methods such as FoldX can serve as reliable initial filters, prioritizing substitutions that experiments would likely confirm as neutral or enhancing. Subsequent application of machine-learning models—ideally fine-tuned on expanded, standardized datasets—could then refine the ranking by capturing higher-order or sequence-contextual effects.

To support these hybrid workflows, future data-generation efforts should mirror AbDesign’s rigor. Incorporating complementary measures, such as SPR-derived kinetics (k_on_, k_off_) and thermostability assays, would provide richer biophysical annotations. Expanding mutational coverage to include double or triple variants, while maintaining uniform protocols, would also illuminate epistatic interactions critical to affinity maturation. Finally, establishing community-wide blind-prediction challenges on withheld AbDesign subsets would encourage transparent, comparable assessments and drive method improvement.

In conclusion, AbDesign demonstrates that a moderately sized but meticulously curated dataset can reveal the current limits of computational antibody-affinity prediction and reaffirm the enduring value of physics-based calculations. As the field moves toward truly de novo antibody design, the synergy of expanded, standardized data, hybrid modelling frameworks, and rigorous benchmarking will be essential to transform in silico promise into clinical reality. We make AbDesign available to the community.

## Acknowledgements

The authors thank Lasse Møller Blaabjerg for his valuable scientific discussions and for reviewing a draft of this manuscript.

## Notes

https://naturalantibody.com/ab-design/

### Competing Interest Statement

The authors have declared no competing interest.

## References

1. Crescioli, S. et al. Antibodies to watch in 2024. MAbs 16, 2297450 (2024).

2. Wilman, W. et al. Machine-designed biotherapeutics: opportunities, feasibility and advantages of deep learning in computational antibody discovery. Brief. Bioinform. 23, (2022).

3. Bielska, W. et al. Applying computational protein design to therapeutic antibody discovery - current state and perspectives. Front. Immunol. 16, (2025).

4. Bennett, N. R. et al. Atomically accurate de novo design of antibodies with RFdiffusion. Bioengineering (2024).

5. Berman, H. M. et al. The Protein Data Bank. Nucleic Acids Res. 28, 235–242 (2000).

6. Dunbar, J. et al. SAbDab: the structural antibody database. Nucleic Acids Res. 42, D1140–6 (2014).

7. Ferdous, S. & Martin, A. C. R. AbDb: antibody structure database-a database of PDB-derived antibody structures. Database 2018, (2018).

8. Jankauskaite, J., Jiménez-García, B., Dapkunas, J., Fernández-Recio, J. & Moal, I. H. SKEMPI 2.0: an updated benchmark of changes in protein-protein binding energy, kinetics and thermodynamics upon mutation. Bioinformatics 35, 462–469 (2019).

9. Sirin, S., Apgar, J. R., Bennett, E. M. & Keating, A. E. AB-Bind: Antibody binding mutational database for computational affinity predictions. Protein Sci. 25, 393–409 (2016).

10. Shanehsazzadeh, A. et al. IgDesign: In vitro validated antibody design against multiple therapeutic antigens using inverse folding. Synthetic Biology (2023).

11. Hitawala, F. N. & Gray, J. J. What has AlphaFold3 learned about antibody and nanobody docking, and what remains unsolved? Bioengineering (2024).

12. Mason, D. M. et al. Optimization of therapeutic antibodies by predicting antigen specificity from antibody sequence via deep learning. Nat Biomed Eng 5, 600–612 (2021).

13. Lim, Y. W., Adler, A. S. & Johnson, D. S. Predicting antibody binders and generating synthetic antibodies using deep learning. MAbs 14, 2069075 (2022).

14. Chinery, L. et al. Baselining the Buzz Trastuzumab-HER2 Affinity, and Beyond. bioRxiv 2024.03.26.586756 (2024) doi:10.1101/2024.03.26.586756.

15. Dudzic, P. et al. RIOT-Rapid Immunoglobulin Overview Tool-annotation of nucleotide and amino acid immunoglobulin sequences using an open germline database. Brief. Bioinform. 26, bbae632 (2024).

16. Abanades, B., et al. ImmuneBuilder: Deep-Learning models for predicting the structures of immune proteins. bioRxiv 2022.11.04.514231 (2022) doi:10.1101/2022.11.04.514231.

17. Eastman, P. et al. OpenMM 7: Rapid development of high performance algorithms for molecular dynamics. PLoS Comput. Biol. 13, e1005659 (2017).

18. Hummer, A. M., Schneider, C., Chinery, L. & Deane, C. M. Investigating the volume and diversity of data needed for generalizable antibody-antigen ΔΔG prediction. bioRxiv 2023.05.17.541222 (2023) doi:10.1101/2023.05.17.541222.

19. Jin, W. et al. DSMBind: SE(3) denoising score matching for unsupervised binding energy prediction and nanobody design. Bioinformatics (2023).

20. Shan, S. et al. Deep learning guided optimization of human antibody against SARS-CoV-2 variants with broad neutralization. Proc. Natl. Acad. Sci. U. S. A. 119, e2122954119 (2022).

21. Luo, S. et al. Rotamer Density Estimator is an Unsupervised Learner of the Effect of Mutations on Protein-Protein Interaction. Bioinformatics (2023).

22. Krawczyk, K., Baker, T., Shi, J. & Deane, C. M. Antibody i-Patch prediction of the antibody binding site improves rigid local antibody–antigen docking. Protein Eng. Des. Sel. 26, 621–629 (2013).

23. Birtalan, S. et al. The intrinsic contributions of tyrosine, serine, glycine and arginine to the affinity and specificity of antibodies. J. Mol. Biol. 377, 1518–1528 (2008).

24. Mason, D. M. & Reddy, S. T. Predicting adaptive immune receptor specificities by machine learning is a data generation problem. Cell Syst. 15, 1190–1197 (2024).

25. Bachas, S. et al. Antibody optimization enabled by artificial intelligence predictions of binding affinity and naturalness. bioRxiv 2022.08.16.504181 (2022) doi:10.1101/2022.08.16.504181.

26. Schymkowitz, J. et al. The FoldX web server: an online force field. Nucleic Acids Res. 33, W382–8 (2005).

27. Bennett, N. R. et al. Atomically accurate de novo design of single-domain antibodies. bioRxiv (2024) doi:10.1101/2024.03.14.585103.

28. Lippow, S. M., Wittrup, K. D. & Tidor, B. Computational design of antibody-affinity improvement beyond in vivo maturation. Nat. Biotechnol. 25, 1171–1176 (2007).

29. Gelman, S. et al. Biophysics-based protein language models for protein engineering. bioRxivorg (2025) doi:10.1101/2024.03.15.585128.

