## Supplementary Materials for "AbDesign: Database of point mutants of antibodies with associated structures reveals poor generalization of binding predictions from machine learning models"

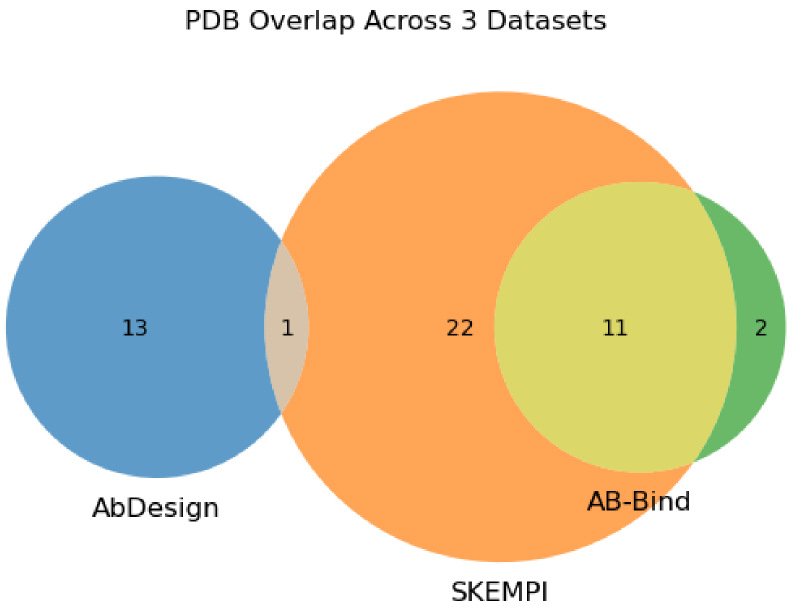


**Supplementary Figure 1. PDB overlap across AbDesign, SKEMPI, and AB-Bind datasets.** SKEMPI shares 11 PDB structures with AB-Bind and 1 with ABDesign, while AbDesign and AB-Bind share no structures. AbDesign is largely non-overlapping, making it a useful independent test set for benchmarking.

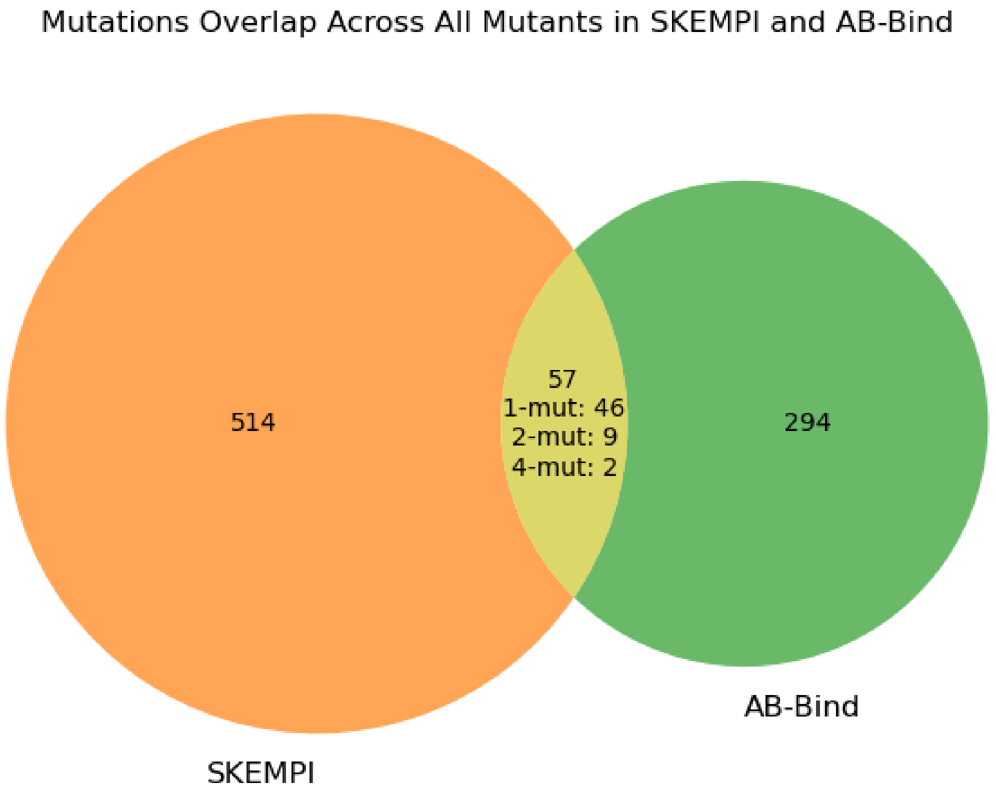


**Supplementary Figure 2. Mutations overlap across all antibody mutants in the SKEMPI and AB-Bind datasets.** A total of 57 mutants are shared between the two datasets, consisting of 46 single-point mutations (*1-mut*), 9 double mutants (*2-mut*), and 2 quadruple mutants (*4-mut*). The majority of mutants in SKEMPI and AB-Bind are unique to each dataset, indicating that they cover different parts of the antibody mutation space.

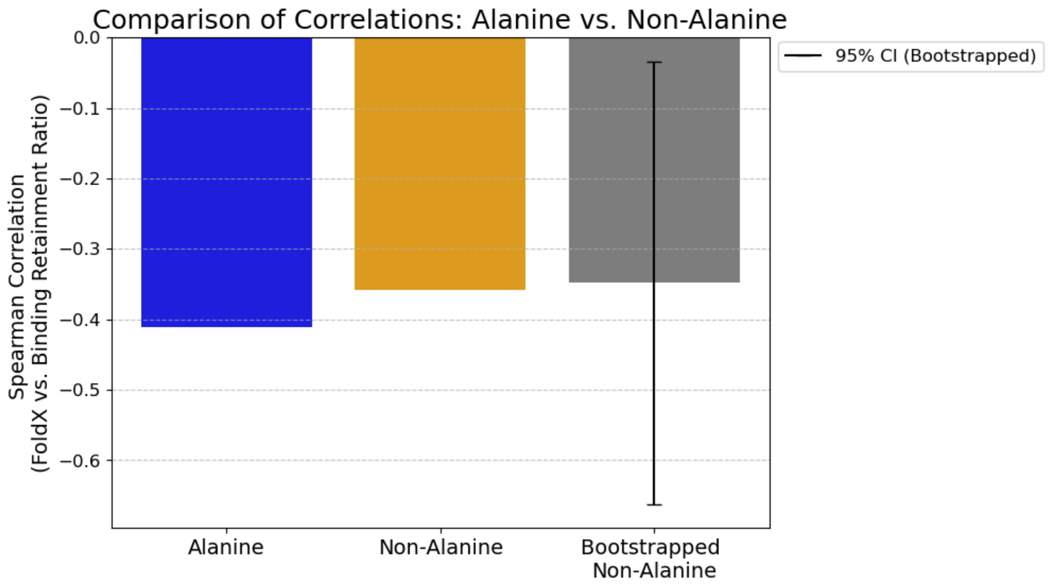


**Supplementary Figure 3. Alanine vs. Non-Alanine correlations to FoldX.** This figure compares FoldX $\Delta\Delta G$ to the binding retainment ratio, therefore a negative correlation is expected. We observe  a drop in correlation for Non-Alanine mutations, despite FoldX being a biophysical method that does not rely on biased training data, although the difference is not statistically significant. Due to the imbalance between Alanine and Non-Alanine mutations (36 vs. 391), bootstrapping was applied to the Non-Alanine group, resampling the data 10,000 times. The bootstrapped correlation of (-0.349) is close to the observed Non-Alanine correlation (-0.358). The 95% confidence interval (CI) [-0.647, -0.003] is wide, indicating high variability in the correlation estimates, but confirms a negative correlation.

**Supplementary Table 1. Means of PDB-stratified correlations.** For each PDB separately in every dataset-methods-structure type cohort, we calculated the correlation between model scores and experimental values. Those correlations were average thereafter to produce a single measure. Note: there are multiple point mutations in SKEMPI and AB-Bind included in these results.

| DDG predictor - Structure prediction method | Dataset | PDB Stratified Spearman Mean Correlation | PDB Stratified Pearson Mean Correlation |
|---|---|---|---|
| DSMBind conformation | SKEMPI (train only) | 0.27 | 0.19 |
| DSMBind conformation | SKEMPI (full) | 0.27 | 0.16 |
| DSMBind conformation | AB-Bind | 0.16 | 0.17 |
| DSMBind conformation | AbDesign | 0.06 | 0.04 |
| DSMBind abb2 | SKEMPI (train only) | 0.45 | 0.36 |
| DSMBind abb2 | SKEMPI (full) | 0.37 | 0.35 |
| DSMBind abb2 | AB-Bind | 0.24 | 0.25 |
| DSMBind abb2 | AbDesign | -0.11 | -0.04 |
| binding-ddg-pred conformation | SKEMPI (train only) | 0.74 | 0.75 |
| binding-ddg-pred conformation | SKEMPI (full) | 0.67 | 0.72 |
| binding-ddg-pred conformation | AB-Bind | 0.62 | 0.50 |
| binding-ddg-pred conformation | AbDesign | 0.11 | 0.13 |
| binding-ddg-pred abb2 | SKEMPI (train only) | 0.49 | 0.55 |
| binding-ddg-pred abb2 | SKEMPI (full) | 0.44 | 0.50 |
| binding-ddg-pred abb2 | AB-Bind | 0.55 | 0.45 |
| binding-ddg-pred abb2 | AbDesign | 0.18 | 0.20 |
| RDE-PPI wt | SKEMPI (train only) | 0.99 | 0.99 |

| | | | |
|---|---|---|---|
| RDE-PPI wt | SKEMPI (full) | 0.91 | 0.93 |

**Supplementary Table 2. Total correlations, without PDB-stratification.** Correlations for all PDBs were pooled for each method. Note: there are multiple point mutations in SKEMPI and AB-Bind included in these results.

| DDG predictor - Structure prediction method | Dataset | Spearman Correlation | Pearson Correlation |
|---|---|---|---|
| DSMBind conformation | SKEMPI (train only) | 0.27 | 0.19 |
| DSMBind conformation | SKEMPI (full) | 0.27 | 0.16 |
| DSMBind conformation | AB-Bind | 0.16 | 0.17 |
| DSMBind conformation | AbDesign | 0.06 | 0.04 |
| DSMBind abb2 | SKEMPI (train only) | 0.45 | 0.36 |
| DSMBind abb2 | SKEMPI (full) | 0.37 | 0.35 |
| DSMBind abb2 | AB-Bind | 0.24 | 0.25 |
| DSMBind abb2 | AbDesign | -0.11 | -0.04 |
| binding-ddg-pred conformation | SKEMPI (train only) | 0.74 | 0.75 |
| binding-ddg-pred conformation | SKEMPI (full) | 0.67 | 0.72 |
| binding-ddg-pred conformation | AB-Bind | 0.62 | 0.50 |
| binding-ddg-pred conformation | AbDesign | 0.11 | 0.13 |
| binding-ddg-pred abb2 | SKEMPI (train only) | 0.49 | 0.55 |
| binding-ddg-pred abb2 | SKEMPI (full) | 0.44 | 0.50 |
| binding-ddg-pred abb2 | AB-Bind | 0.55 | 0.45 |
| binding-ddg-pred abb2 | AbDesign | 0.18 | 0.20 |

| | | | |
|---|---|---|---|
| RDE-PPI wt | SKEMPI (train only) | 0.99 | 0.99 |
| RDE-PPI wt | SKEMPI (full) | 0.91 | 0.93 |
| RDE-PPI wt | AB-Bind | 0.58 | 0.54 |
| RDE-PPI wt | AbDesign | -0.07 | -0.07 |
| RDE-PPI wt_abb2 | SKEMPI (train only) | 0.92 | 0.92 |
| RDE-PPI wt_abb2 | SKEMPI (full) | 0.86 | 0.87 |
| RDE-PPI wt_abb2 | AB-Bind | 0.6 | 0.55 |
| RDE-PPI wt_abb2 | AbDesign | 0.11 | 0.15 |

**Supplementary Table 3. Binding energy prediction-measurement correlations for alanine mutations only.** We contrasted the predictive ability of the models to assess the effects of alanine mutations on binding energy. Pearson's correlations were calculated on mutations to alanine or to any other amino acid. Correlations were obtained by either pooling all the predictions to achieve a single correlation or computing on a per-PDB basis with the mean reported. We note the better prediction in bold. In cases where it is ambiguous to call which outcome is superior (e.g. nil versus negative correlation), we left results unmarked.

| DDG predictor - Structure prediction method | Dataset | Unstratified Pearson Correlation | | PDB Stratified Pearson Mean Correlation | |
|---|---|---|---|---|---|
| | | Non-alanine | Alanine | Non-alanine | Alanine |
| DSMBind conformation | SKEMPI (train only) | 0.17 | 0.17 | 0.15 | 0.06 |
| DSMBind conformation | SKEMPI (full) | 0.18 | 0.19 | 0.23 | -0.12 |
| DSMBind conformation | AB-Bind | 0.09 | 0.13 | -0.03 | 0.36 |
| DSMBind conformation | AbDesign | 0.03 | 0.16 | 0.00 | 0.26 |
| DSMBind abb2 | SKEMPI (train only) | 0.32 | 0.17 | 0.20 | 0.10 |
| DSMBind abb2 | SKEMPI (full) | 0.28 | 0.22 | 0.24 | -0.04 |
| DSMBind abb2 | AB-Bind | 0.11 | 0.16 | 0.00 | 0.20 |
| DSMBind abb2 | AbDesign | -0.03 | -0.08 | 0.01 | 0.12 |
| binding-ddg-pred conformation | SKEMPI (train only) | 0.68 | 0.76 | 0.77 | 0.35 |
| binding-ddg-pred conformation | SKEMPI (full) | 0.69 | 0.81 | 0.60 | 0.52 |
| binding-ddg-pred conformation | AB-Bind | 0.27 | 0.65 | 0.44 | 0.44 |
| binding-ddg-pred conformation | AbDesign | 0.13 | 0.23 | 0.09 | -0.01 |

| | | | | | |
|---|---|---|---|---|---|
| binding-ddg-pred abb2 | SKEMPI (train only) | 0.55 | 0.65 | 0.56 | 0.28 |
| binding-ddg-pred abb2 | SKEMPI (full) | 0.57 | 0.69 | 0.49 | 0.39 |
| binding-ddg-pred abb2 | AB-Bind | 0.29 | 0.48 | 0.18 | 0.42 |
| binding-ddg-pred abb2 | AbDesign | 0.19 | 0.4 | 0.13 | 0.34 |
| RDE-PPI wt | SKEMPI (train only) | 0.99 | 0.99 | 0.97 | 0.97 |
| RDE-PPI wt | SKEMPI (full) | 0.89 | 0.97 | 0.83 | 0.98 |
| RDE-PPI wt | AB-Bind | 0.38 | 0.75 | 0.73 | 0.81 |
| RDE-PPI wt | AbDesign | -0.07 | -0.12 | 0.01 | 0.09 |
| RDE-PPI wt_abb2 | SKEMPI (train only) | 0.92 | 0.89 | 0.80 | 0.84 |
| RDE-PPI wt_abb2 | SKEMPI (full) | 0.84 | 0.88 | 0.69 | 0.82 |
| RDE-PPI wt_abb2 | AB-Bind | 0.40 | 0.72 | 0.40 | 0.79 |
| RDE-PPI wt_abb2 | AbDesign | 0.15 | 0.3 | 0.18 | 0.22 |

**Supplementary Table 4. Categorization of common biophysical properties of amino acids.**

| Biochemical Categories | Amino Acids | Size Categories | Amino Acids |
|---|---|---|---|
| Acidic | D, E | Aromatic | F, Y, W |
| Basic | K, R, H | Large | M, I, L, K, R |
| Hydrophobic | A, V, L, I, M, F, W, Y | Medium | P, D, N, T, E, Q, H, V |
| Polar | S, T, N, Q | Small | S, C |
| Special | C, G, P | Tiny | G, A |

**Supplementary Table 5. Predictive performance on the reduced FoldX dataset.** For 9 out of 14 PDBs in our AbDesign dataset, we could obtain FoldX predictions. For each individual PDB, we show the Spearman's correlation coefficient between the model score and the binding retainment score. Mean or max correlations across structure prediction approaches are shown for each ML model. As FoldX uses its own structure prediction approach, leading to a single prediction, the mean and max correlations are identical.

| PDB ID | FoldX | binding-ddg-pred | | DSMBind | | RDE-PPI | |
|---|---|---|---|---|---|---|---|
| | | max | mean | max | mean | max | mean |
| 1bj1 | 0.29 | -0.18 | -0.18 | -0.40 | -0.45 | 0.13 | 0.02 |
| 5f9w | 0.26 | 0.09 | 0.06 | 0.01 | -0.01 | -0.01 | -0.03 |
| 5ggv | 0.52 | 0.28 | 0.24 | -0.26 | -0.35 | -0.29 | -0.37 |
| 6mfp | 0.35 | -0.11 | -0.13 | 0.06 | 0.00 | -0.11 | -0.20 |
| 6nms | 0.83 | 0.17 | 0.12 | 0.23 | 0.23 | 0.14 | -0.07 |
| 6xkr | 0.08 | -0.15 | -0.16 | -0.07 | -0.08 | -0.14 | -0.16 |
| 7bej | 0.37 | -0.37 | -0.42 | -0.06 | -0.10 | 0.18 | -0.07 |
| 7jmo | 0.45 | -0.08 | -0.17 | 0.16 | 0.13 | 0.14 | 0.03 |
| 7kf0 | 0.70 | -0.42 | -0.57 | 0.16 | 0.08 | -0.16 | -0.35 |
